# Synergistic CRISPR-Cas Antimicrobials through Essential and Defensive Gene Cotargeting in *Staphylococcus aureus*

**DOI:** 10.64898/2026.06.25.734632

**Authors:** David S. Dooley, Cong T. Trinh

## Abstract

Multidrug-resistant pathogens pose a major threat to One Health. Within the past decade, CRISPR-Cas systems have been explored as sequence-specific antimicrobials. While chromosomal injury has been considered the primary mechanism underlying pathogen killing by CRISPR-Cas antimicrobials, the synergistic role of gene disruption together with chromosomal injuries remains poorly understood. In this study, we characterized a new class of CRISPR-Cas antimicrobials that simultaneously cotarget essential and defensive genes to enhance potency against the clinically relevant pathogen *Staphylococcus aureus*. High-throughput CRISPR screening identified top-performing guide RNAs for twenty functionally diverse essential and defensive genes across the *S. aureus* genome. CRISPR-Cas antimicrobials were modularly formulated to target single or multiple gene loci and packaged in phage-like particles for specific delivery. By engineering an *S. aureus* production host with a chromosomally integrated anti-CRISPR protein, we demonstrated efficient production of CRISPR-Cas antimicrobials targeting any *S. aureus* chromosomal locus without self-targeting. Characterization of CRISPR-Cas antimicrobials with single guide RNA designs revealed that potency varied according to targeted gene function, achieving up to a 4-log_10_ reduction in viability and outperforming traditional antibiotics. Multiplexed configurations were consistently more effective than single-targeting designs, with the top-performing design demonstrating a 4.7-log_10_ reduction in viability. Cotargeting essential and defensive genes revealed synergies that led to improved lethality and attenuated resistance, with enhanced activity in biofilms compared to traditional antibiotics. Genes involved in signaling and stress responses were important defensive targets for developing cotargeting CRISPR-Cas antimicrobials. Overall, this study establishes design principles for synergistic CRISPR-Cas antimicrobials applicable to next-generation precision antimicrobial development.

**SIGNIFICANCE:** The ability to effectively combat multidrug-resistant pathogens is of primary importance to One Health. This study develops a generalizable design principle for formulating potent CRISPR-Cas antimicrobials that exploit synergistic cotargeting strategies for enhanced pathogen killing. In addition to chromosomal injuries, we found that disruption of gene function plays a crucial role in determining the lethality of CRISPR-Cas antimicrobials, providing a generalizable framework for effective CRISPR-Cas antimicrobial design. The development of a CRISPR-Cas antimicrobial production host with stable, chromosomally integrated anti-CRISPR genes greatly expands the modularity, adaptability, and efficiency of formulating CRISPR-Cas antimicrobials and enables deeper insights into the molecular mechanisms involved in eliminating multidrug-resistant pathogens.

## INTRODUCTION

The emergence of antimicrobial-resistant (AMR) pathogens is among the gravest threats to One Health and, according to a recent report, is at least as problematic as HIV or malaria^1^. Unfortunately, since 1987, there has been a “discovery void” of novel classes of antibiotics^2^. Among AMR pathogens, *Staphylococcus aureus* stands out as the fastest-growing cause of AMR-related deaths^3^. Because *S. aureus* is carried by 30% of the population, exposure to this pathobiont is exceedingly common, elevating morbidity in at-risk populations and engendering immunogenic familiarity that confounds effective vaccine development^4,5^. These factors underscore the need to confront virulent, resistant *S. aureus* with innovative therapeutic strategies.

CRISPR-Cas systems hold promise as sequence-specific antimicrobials against *S. aureus*^6^. Originally adaptive immune systems in prokaryotes, CRISPR-Cas systems are RNA-guided endonucleases that target and destroy DNA or RNA substrates with high specificity. When co-opted for use as antimicrobials, they can be programmed to target essential cellular machinery, leading to cell death via nucleic acid degradation and/or protein loss of function. CRISPR-Cas antimicrobials have been successfully developed for several bacterial pathogens, including *Escherichia coli*^7–13^, *S. aureus*^14–19^, *Clostridioides difficile*^20^, and others^21–28^, highlighting their generalizability. These studies, however, have not substantially investigated the effect of targeted gene function on CRISPR-Cas antimicrobial efficiency, instead focusing on chromosomal injury or antibiotic resensitization as a major mode of pathogen neutralization. Recent studies of CRISPR-Cas antifungals suggest that targeted disruption of gene function could enhance killing while minimizing resistance^29^. Therefore, the limited understanding of optimal essential and defensive gene targets and their potential cotargeting synergies represents a significant knowledge gap in the development of CRISPR-Cas antimicrobials against bacterial pathogens such as *S. aureus*.

Specific delivery is critical to precision antimicrobial treatment. Phages can be harnessed as delivery vehicles by packaging CRISPR-Cas antimicrobial cargos on phage-borne DNA^6,14^. Production of CRISPR-carrying phage-like particles (PLPs) requires an *S. aureus* production host, but self-targeting represents a major barrier during this process. The most common strategy to avoid self-targeting is to design guide RNAs (gRNAs) that target a chromosomal locus not present in the production host^14,29,30^. This approach circumvents self-targeting at the cost of heavily restricting target selection. Other solutions include activating endogenous CRISPR systems with exogenous gRNAs^31–33^, using inducible CRISPR systems^32,34^, or employing heterologous production hosts^27,35^; however, these methods also have major limitations. First, target pathogens may not possess endogenous CRISPR machinery, and those that do may have partial or inactive systems. Determination of the protospacer adjacent motif (PAM) also limits this approach, since experimental validation is laborious and *in silico* PAM prediction is unreliable. Second, while inducible systems are a useful test bed for different gRNAs, real-world treatments must act constitutively to achieve high efficacy, safety, and economy. To accurately represent the lethality of CRISPR-Cas antimicrobial formulations *in vitro*, constitutive one-plasmid or phage-embedded systems should be employed. Finally, although phage engineering efforts have greatly advanced in recent years, the ability to rapidly produce phages with tunable host ranges remains elusive, especially for non-model organisms and recently isolated strains, making the utilization of heterologous production hosts highly challenging. Therefore, it is critical to establish a novel production host that can produce self-targeting CRISPR-Cas antimicrobials for any pathogenic locus. Such a production host enables effective antimicrobial formulation and advances understanding of the molecular mechanisms of gene target selection and action for therapeutic deployment.

In this study, we investigated the synergistic cotargeting of essential and defensive genes to elucidate how the functional roles of gene targets impact the potency of CRISPR-Cas antimicrobials beyond chromosomal injury. We started by screening a library of gRNAs against twenty functionally diverse essential and defensive genes across the *S. aureus* genome to assess their activities, with four guides targeting each gene. To evaluate the top-performing guides for CRISPR-Cas antimicrobials in isolation and combination using phage delivery, we engineered an *S. aureus* production strain that harbors an SpCas9 anti-CRISPR to avoid self-targeting. Top-performing guides for each gene were evaluated individually before being combinatorially cotargeted to evaluate synergistic interactions.

## RESULTS AND DISCUSSION

### A pooled CRISPR library screen identifies highly active guides targeting essential and defensive genes in *S. aureus*

We developed a pooled CRISPR library screen to identify highly active guides for CRISPR-Cas antimicrobial formulation in *S. aureus* (**Fig. 1A**). CRISPR-Cas antimicrobials are modularly designed, comprising a Cas submodule, a gRNA submodule, and a phage packaging module (**Fig. 1B**). As a proof-of-study, we designed gRNAs to target ten essential and ten defensive genes in *S. aureus* with four guides per gene using the gRNA design software, CASPER^36^. We chose a set of ten essential genes by filtering the Database of Essential Genes^37^ for genes conserved across all strains of *S. aureus*, separating genes by functional category with Clusters of Orthologous Genes^38^ (COG) annotations that span replication (*dnaA, dnaJ*), transcription (*rpoB*, *nusA*), translation (*rpsL*, *trmD*, *rpmD*), metabolism (*nadE*, *eno, secY*), and protein trafficking (*secY*), and selecting genes from each COG (**Fig. 1C**). We selected a set of defensive genes based on literature evidence with a focus on DNA repair (*umuC*, *cinA*, *recA*, *ligA, recF*), stress response (*clpP, dnaK*), and cell signaling (*sosA*, *lexA*, *sigB*) (**Fig. 1C**). As controls, we included two guides from literature^39^, two multitargeting, two non-coding, and two null-targeting (NT) guides, bringing the library to a total size of 88.

**Figure 1:**
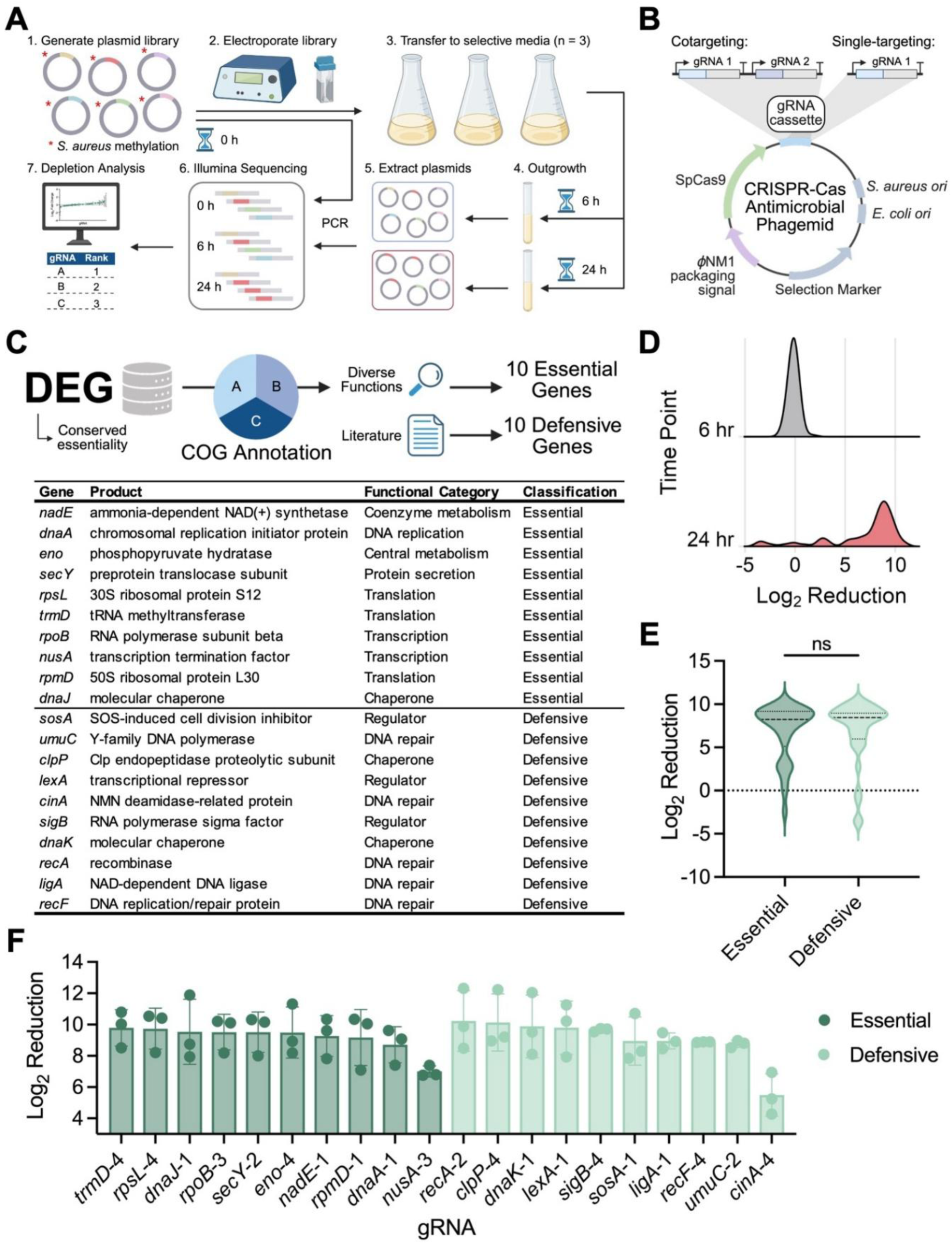
High-throughput pooled CRISPR library screen reveals highly active gRNAs against *S. aureus* essential and defensive genes. (**A**) Experimental workflow for CRISPR screen. (**B**) Schematic of the modular design of CRISPR-Cas antimicrobials. (**C**) Workflow and results for essential and defensive gene selection. (**D**) Distribution of gRNA log_2_ reduction values at 6 hr and 24 hr time points. (**E**) Comparison of log_2_ reduction values between essential and defensive gene categories. Statistical significance determined by unpaired t test. (**F**) Log_2_ reduction values for the top-performing guide of each gene.

Before transformation into *S. aureus* Newman, we passaged the guide library through *E. coli* IM08*β* to confer the correct methylation profile for avoiding restriction^40^. Sequencing of pre-transformed libraries revealed that nine guides were lost during the cloning process, and a further five guides were lost during passage through IM08*β*, resulting in a final library size of 74 (**Supplementary Fig. S1A)**. Despite these losses, all 20 target genes were represented by at least two gRNAs. After transformation, all sequencing samples showed high gRNA coverage (*>*3000x) and mapping rate (*>*80%) (**Supplementary Figs. S1B, S1C**). Importantly, no gRNAs were lost between 0 hr and 6 hr time points, indicating that transformation efficiency was not limiting (**Supplementary Fig. S1A**). Log_2_ reductions between the pre-transformed and 6 hr post-transformed libraries showed minimal changes, with a median of -0.117, indicating that 6 hr was not enough time to significantly perturb the gRNA population (**Figs. 1D, S1D, S1E**). Therefore, the 0 hr time point was used as the reference for guide depletion. By 24 hr, most guides were heavily depleted, with a median log_2_ reduction of 8.13, underscoring the sensitivity of *S. aureus* to CRISPR-Cas activities that cause chromosomal DNA damage and gene disruption (**Figs. 1D, 1F, S1D, S1F**). When comparing the mean log_2_ reduction between essential gene-targeting and defensive gene-targeting guides, we observed no significant difference (**Fig. 1F**). This lack of separation is consistent with the near-saturation depletion of guides at 24 hr post-transformation (**Supplementary Figs. S1D, S1F**), suggesting that chromosomal injury obfuscates functional contributions over long-term exposure to gRNAs. To validate the library results and enable better discrimination between gRNA target groups, we next sought to characterize individual gRNAs *in vitro*.

### An integrated anti-CRISPR protein enables efficient production of self-targeting CRISPR-Cas antimicrobials in *S. aureus*

The generation of PLPs harboring CRISPR-Cas antimicrobials for delivery has long faced a significant barrier due to gRNA self-targeting, which limits the antimicrobial design space^14,30^. Generation of CRISPR-Cas PLPs begins with transformation of the therapeutic phagemid into the production strain. Electroporation of an NT CRISPR phagemid into a standard laboratory strain lysogenized with the *ϕ*NM1 prophage (*S. aureus* SaDD0001) yielded 1.96 ± 1.18×10^5^ transformants/µg using a standard electroporation protocol^41^. In contrast, when a self-targeting gRNA was introduced, not even 1 µg of purified phagemid DNA was able to generate a transformant. Presumably, even if transformants could be obtained, the likelihood of mutation in the CRISPR system would be high. To address these shortcomings, we developed a novel, CRISPR-resistant *S. aureus* production strain (*S. aureus* SaDD0006) to produce self-targeting PLPs. Using traditional allelic exchange, we stably integrated AcrIIA4, an SpCas9 anti-CRISPR protein^42^, at the intergenic site between KMZ21_RS00180 and KMZ21_RS00185 in *S. aureus* SaDD0001 under the control of the strong constitutive promoter SarA P1 and flanked by the lambda t0 terminator (**Fig. 2A**). Presence of AcrIIA4 was verified using PCR and sequencing, revealing correct insertion at the desired locus (**Fig. 2B**).

**Figure 2:**
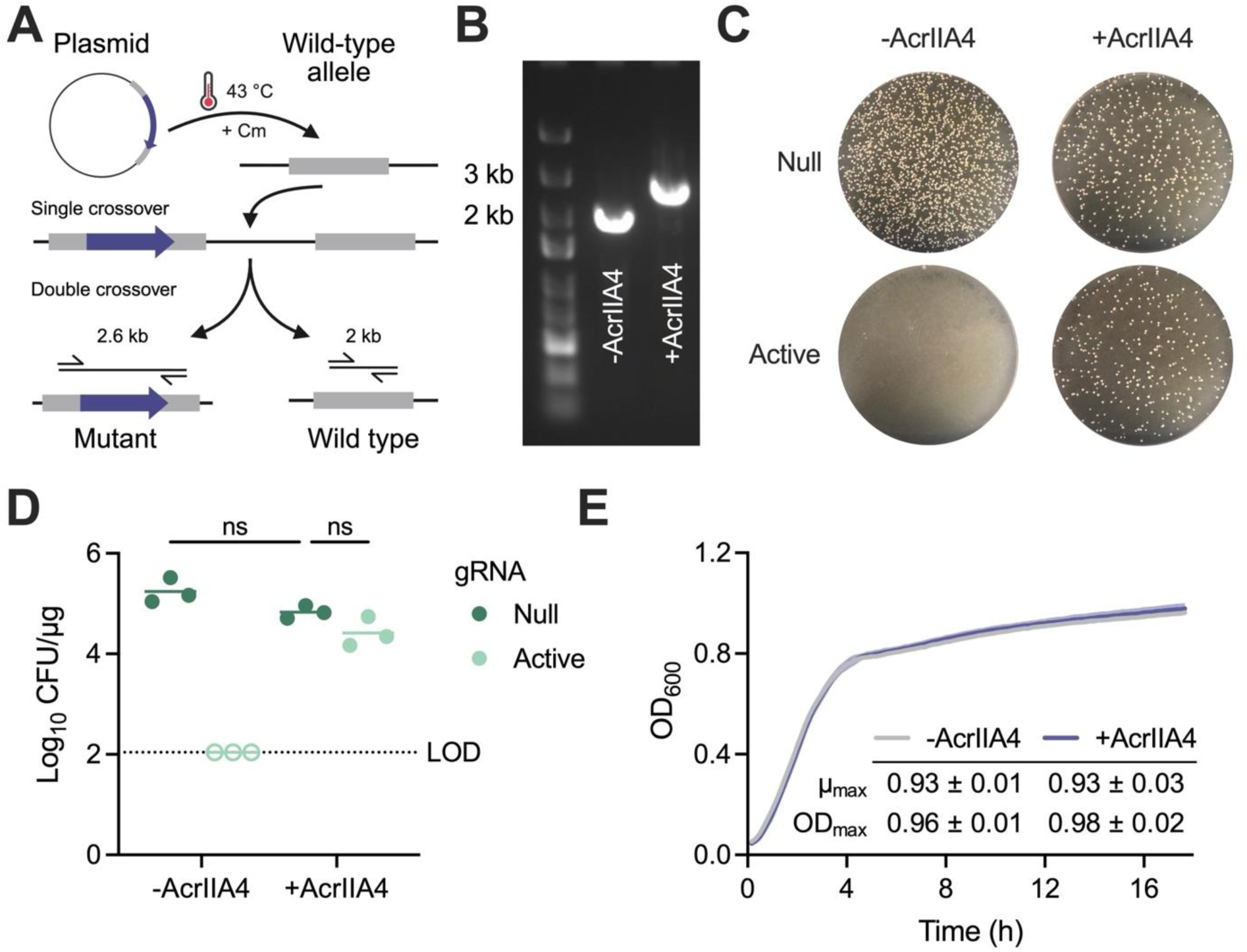
Generation of an anti-CRISPR production strain of *S. aureus* for producing CRISPR-Cas antimicrobials without self-targeting. (**A**) Allelic exchange workflow for inserting AcrIIA4 into *S. aureus* SaDD0001. (**B**) DNA gel of PCR confirming AcrIIA4 knock-in. (**C**) Representative transformations of 100 ng of either null or active CRISPR phagemids into *S. aureus* with and without AcrIIA4. (**D**) Transformation efficiencies of null and active CRISPR phagemids into *S. aureus* with and without AcrIIA4. Open circles indicate measurements below the limit of detection (LOD). Statistical significance determined by two-way ANOVA. (**E**) Growth profiles and metrics of *S. aureus* with and without AcrIIA4.

To test the ability of AcrIIA4 to protect *S. aureus* from CRISPR self-targeting, phagemids with constitutive SpCas9 and gRNA expression were electroporated into strains SaDD0001 and SaDD0006 (**Fig. 2C)**. Phagemids expressing a null guide transformed into both strains with high efficiency, while phagemids with an active guide only transformed into SaDD0006 (**Figs. 2C, 2D**). There was no significant difference between the transformation efficiency of null and active phagemids in SaDD0006, indicating AcrIIA4 provided robust protection from CRISPR self-targeting (**Fig. 2D**). Additionally, the production strain showed no defect in growth rate or carrying capacity compared to the parent strain, indicating that expression of AcrIIA4 imposes minimal metabolic burden on *S. aureus* (**Fig. 2E**). *S. aureus* SaDD0006 was subsequently used to produce CRISPR-Cas PLPs for *in vitro* characterization (**Supplementary Fig. S2**).

Overall, we successfully developed an engineered anti-CRISPR *S. aureus* production strain that enables the generation of CRISPR-Cas antimicrobials targeting any locus in *S. aureus*. This development overcomes current limitations in engineering CRISPR-Cas antimicrobials and expands their design space with high degrees of modularity, adaptability, and efficiency, a capability that is critical for combating rapidly evolving and resistant *S. aureus*.

### Combined chromosomal injury and gene disruption contribute to the potency of CRISPR-Cas antimicrobials

CRISPR-Cas9 systems are known to create blunt end cuts, causing chromosomal injury that induces cell death^43^. To elucidate whether disruption of gene functions contributes to the potency of CRISPR-Cas antimicrobials, we performed individual characterization of top-performing gRNAs targeting either essential or defensive genes (**Fig. 1F**). The most depleted guide for each gene was individually cloned into a phagemid with constitutive SpCas9 and gRNA expression for detailed characterization (**Fig. 1B**). We infected exponentially grown SaDD0001 cells with purified PLPs harboring these antimicrobials at a multiplicity of infection (MOI) of one and determined their potency by comparing the surviving colonies to an NT control (**Fig. 3A**). CRISPR-Cas antimicrobials eliminated *S. aureus* SaDD0001 with varying degrees of efficiency, with log_10_ reductions spanning from 2.82 to 4.05 (**Figs. 3B, 3C**).

**Figure 3:**
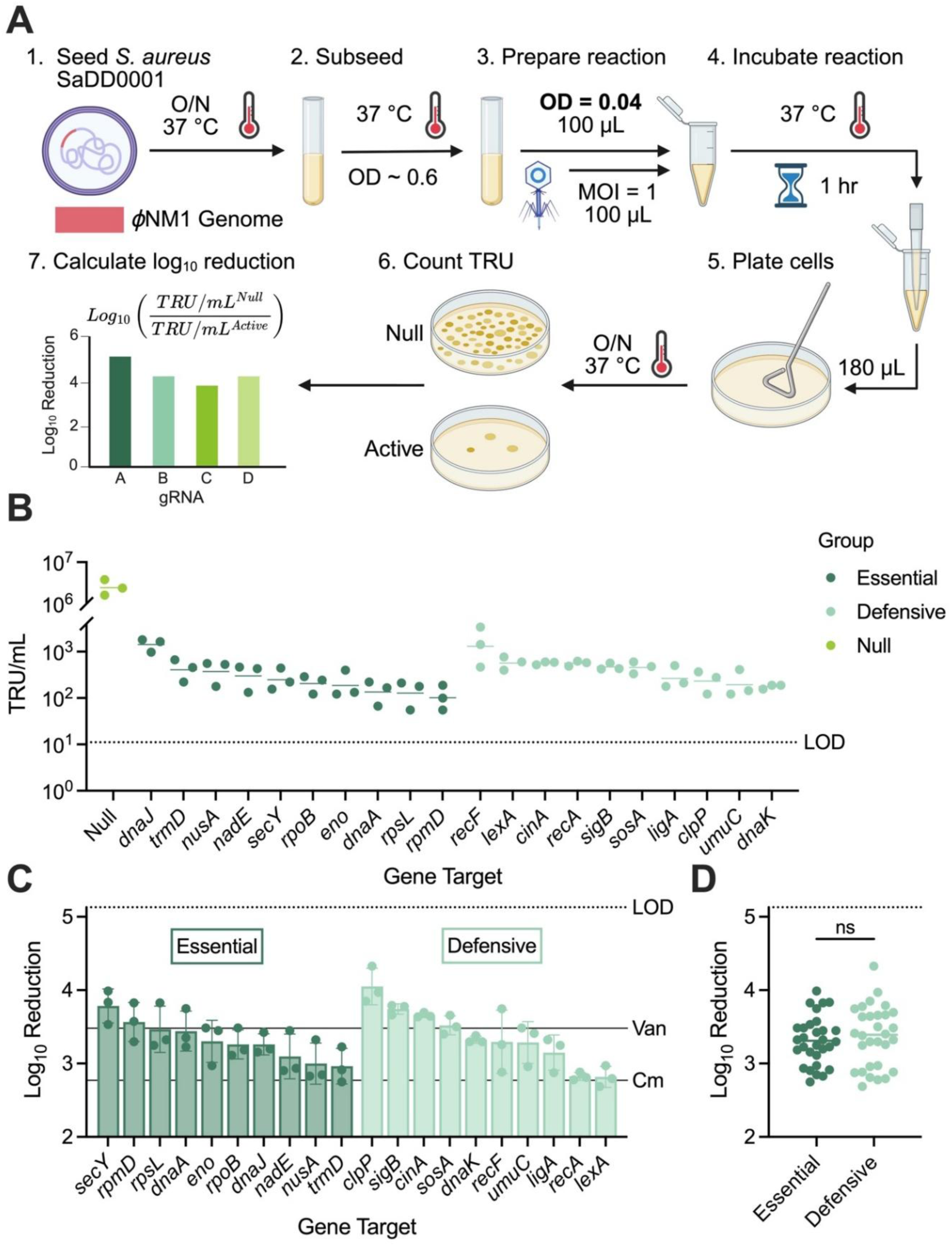
Evaluation of top-performing gRNAs targeting ten essential and ten defensive genes in *S. aureus*. (**A**) Workflow of transduction-based killing assay for evaluating CRISPR-Cas antimicrobials packaged in PLPs. (**B**) Transducing titer of PLPs harboring single gRNAs. Note: Null data is representative. (**C**) Log_10_ reduction values of single gRNAs relative to a null control. Van = vancomycin, Cm = chloramphenicol. (**D**) Comparison of log_10_ reduction values between essential and defensive genes. Statistical significance determined by unpaired t test. LOD = limit of detection.

For CRISPR-Cas antimicrobials targeting single essential genes, we found that the top three lethal essential gene targets were *secY*, *rpmD*, and *rpsL* with log_10_ reductions of 3.79 ± 0.23, 3.57 ± 0.27, and 3.47 ± 0.32, respectively (**Fig. 3C**). *SecY* encodes a protein translocase subunit and *rpmD* and *rpsL* encode ribosomal proteins. In contrast, the worst-performing essential gene targets were *trmD*, *nusA*, and *nadE*, with log_10_ reductions of 2.96 ± 0.24, 3.00 ± 0.28, and 3.10 ± 0.31, respectively. *TrmD* encodes a tRNA methyltransferase, *nusA* encodes a transcription elongation factor, and *nadE* encodes a nicotinamide adenine dinucleotide (NAD) synthetase (**Fig. 3C**).

For CRISPR-Cas antimicrobials targeting single defensive genes, we identified the top three defensive gene targets to be *clpP*, *sigB*, and *cinA* with log_10_ reductions of 4.05 ± 0.25, 3.74 ± 0.07, and 3.66 ± 0.03, respectively (**Fig. 3C**). *ClpP* encodes a stress-responsive protease, *sigB* encodes a master stress response regulator, and *cinA* encodes a nicotinamide mononucleotide deamidase that recycles NAD precursors. The worst-performing defensive gene targets were *lexA*, *recA*, and *ligA*, with log_10_ reductions of 2.82 ± 0.14, 2.82 ± 0.05, and 3.15 ± 0.24, respectively. *LexA* encodes a master transcriptional repressor that regulates the DNA damage-induced SOS response in *S. aureus*, *recA* encodes a recombinase that mediates recombinational DNA repair, and *ligA* encodes a DNA ligase that seals single-strand DNA breaks (**Fig. 3C**).

The observed target-specific variability in lethality indicates that gene functions contribute to CRISPR-Cas antimicrobials beyond chromosomal injuries. Protein biogenesis (e.g., *rpmD* and *rpsL*) and proteostasis (e.g., *secY* and *clpP*) are functional nodes that appear particularly sensitive to CRISPR-mediated destruction. Surprisingly, proteins explicitly involved in DNA repair, such as *recA*, *ligA*, *umuC*, and *lexA*, were not among the top-performing defensive gene targets, suggesting DNA repair may already be limiting before functional knockdown. This conclusion is partially supported by the fact that *S. aureus* has a small *lexA* regulon of only 16 genes, compared to the more robust 43- and 64-gene regulons of *E. coli* and *Bacillus subtilis*, respectively^44^. As such, general stress response pathways (e.g., *cinA*, *sigB*) may be more beneficial to target in *S. aureus*. At a high level, we found that the mean log_10_ reductions for targeting essential genes (3.31 ± 0.33) and defensive genes (3.37 ± 0.41) were similar (**Fig. 3D**), suggesting that chromosomal damage is dominant in *S. aureus* and that gene disruption provides an additional but comparable contribution.

### Synergistic gene cotargeting enhances the potency of CRISPR-Cas antimicrobials

Next, we investigated whether cotargeting essential and defensive genes (E+D) could improve the lethality of CRISPR-Cas antimicrobials over single essential (E), single defensive (D), essential-essential (E+E), or defensive-defensive (D+D) formulations, as recently demonstrated in *S. cerevisiae*^31^. Using killing efficiencies from the individual gRNA characterization (**Fig. 3C**), an essential- and defensive-targeting guide was chosen from each of three regimes—high efficiency (H), medium efficiency (M), and low efficiency (L)—to capture synergistic effects between gRNAs of varying activities. Cotargets were named according to their gene target (E or D) and efficiency (H, M, or L). For example, the HE+MD cotarget combined a high-efficiency essential guide with a medium-efficiency defensive guide. The selected guides were *secY* -2 (HE), *rpoB*-3 (ME), *trmD*-4 (LE), *clpP*-4 (HD), *dnaK*-1 (MD), and *lexA*-1 (LD). Guide pairs were combined in a position-independent manner since no difference in activity was observed when an essential gRNA was paired with a null guide in the first or second position of the cotargeting construct **(Supplementary Fig. S3**). Each gRNA was also paired with itself (“self” constructs) to control for effects arising from differing gRNA expression between single-targeting and cotargeting constructs.

Cotarget phagemids were transformed into *S. aureus* SaDD0006, and PLPs were generated, concentrated, and characterized as before. Cotargets exhibited a wide range of lethalities, with log_10_ reductions of bacterial viability ranging from 3.70 to 4.73 (**Figs. 4A, 4B**). The top five most lethal cotargets were HD+MD (4.73 ± 0.21), HE+ME (4.73 ± 0.16), ME+LD (4.72 ± 0.17), HD+ME (4.65 ± 0.15), and ME+MD (4.63 ± 0.26). The five least lethal cotargets were LE+LD (3.73 ± 0.28), MD+LD (3.93 ± 0.21), HE+LD (4.00 ± 0.16), ME+LE (4.14 ± 0.06), and MD+LE (4.24 ± 0.26). All the top-performing cotargets included either ME (*rpoB*-3) or MD (*dnaK*-1), and all the worst-performing cotargets included either LE (*trmD*-4) or LD (*lexA*-1).

**Figure 4:**
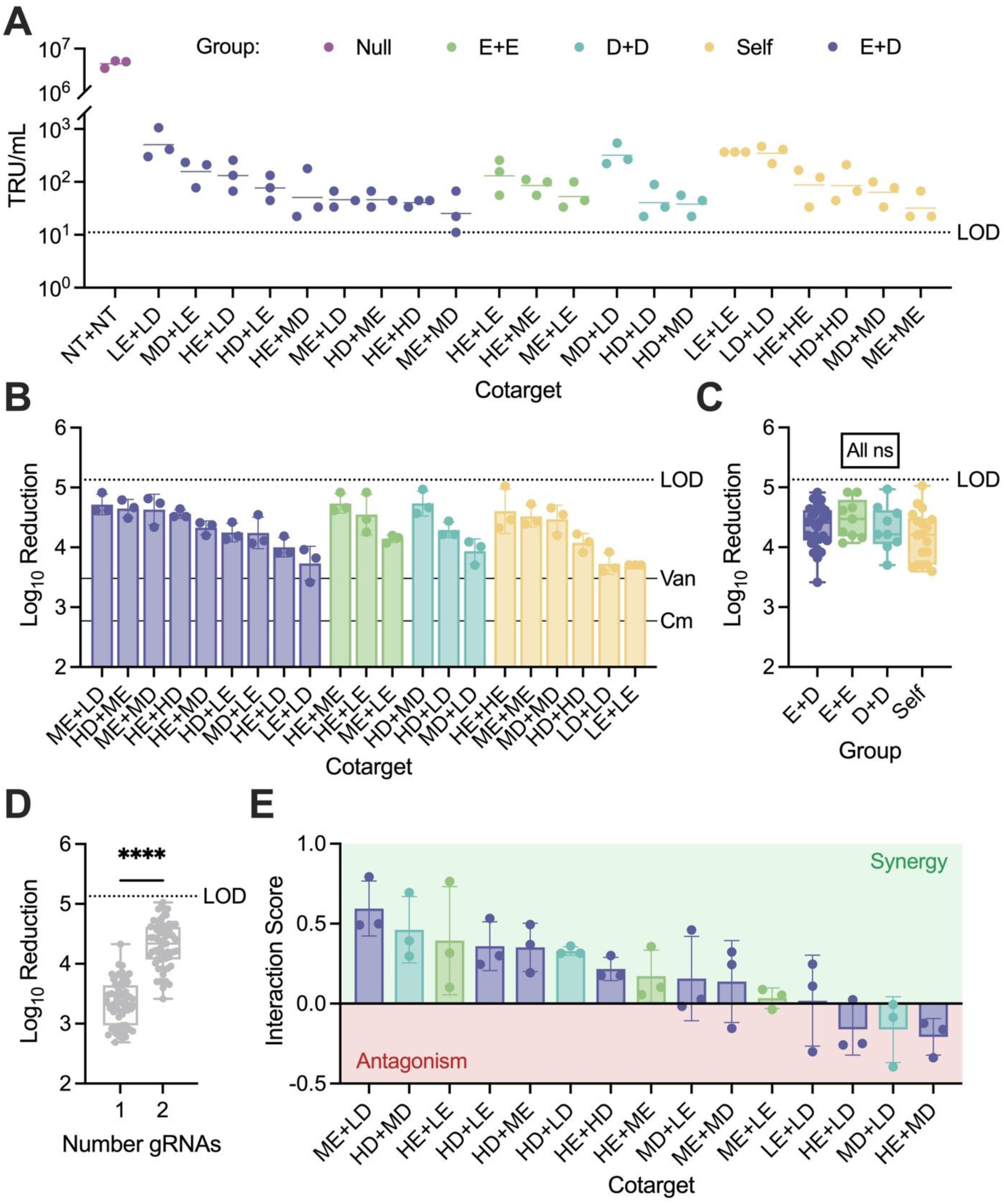
Evaluation of essential and defensive gene cotargeting in *S. aureus*. (**A**) Transducing titer of PLPs for all cotargets. (**B**) Log_10_ reduction values of cotargets relative to a null control. Van = vancomycin, Cm = chloramphenicol. (**C**) Summary of log_10_ reduction values according to cotargeting groups. (**D**) Summary of log_10_ reduction values according to gRNA number. (**E**) Interaction score analysis of cotargets. Statistical significance determined by (**C**) one-way ANOVA and (**D**) unpaired t test. LOD = limit of detection.

There were no significant differences in the lethality of constructs according to cotargeting group (**Fig. 4C**), but in general, cotargeting was more effective than single-targeting at eliminating *S. aureus*—4.31 ± 0.38 vs. 3.34 ± 0.37 log_10_ reduction, respectively (**Fig. 4D**). Even self constructs outperformed cognate single-targeting constructs by an average of 0.82 logs, indicating that SpCas9 is not limiting in our experimental design. Although two copies were more effective, the relative activities of the guide RNAs remained generally consistent between self and single constructs. Both LE+LE and LD+LD constructs were again the worst performers and HE+HE was a top performer. HD+HD, however, did not largely benefit from a second copy of the gRNA, falling to the fourth most active self construct.

To evaluate the synergy of guide pairs, interaction score analysis was performed (**Fig. 4E**). The most synergistic guide pair was an essential and defensive cotarget of ME (*rpoB*-3) and LD (*lexA*-1), which outperformed the expected log_10_ reduction by 0.59 ± 0.17 orders of magnitude and was one of the most lethal designs overall. In *Clostridium thermocellum*, *rpoB* is part of the LexA regulon^45^, suggesting RNA polymerase expression is heightened under stressful conditions to accommodate increased transcription of SOS response effectors. Therefore, we hypothesize that cotargeting *rpoB* and *lexA* launches a multi-pronged attack at critical nodes of the SOS response, collapsing the entire network in a way single-targeting cannot. Notably, neither ME nor LD gRNAs were among the most lethal single-targeting designs, suggesting that effective cotargets cannot be predicted solely from individual guide activities and highlighting the significance of gene function in determining the potency of CRISPR-Cas antimicrobials.

The next most synergistic cotarget was HD (*clpP*-4) with MD (*dnaK*-1), with an interaction score of 0.46 ± 0.21. In *S. aureus*, the loss of ClpP results in the accumulation of misfolded proteins, which in turn upregulates the expression of chaperones and proteases like DnaK^46^. ClpP and DnaK have also been shown to physically interact in *S. aureus* and *E. coli*^47,48^, strengthening the functional link between these two proteins. As with ME+LD, simultaneous targeting of *clpP* and *dnaK* compromises multiple nodes of an essential cellular process—in this case, protein homeostasis and SOS response. The strength of this approach is complemented by the fact that *S. aureus* deletion mutants of *clpP* or *dnaK* have reduced fitness and/or attenuated virulence, which may force escapee cells into an evolutionary trap. Based on our nomenclature, HD+MD is a defensive-defensive cotarget, but in some strains of *S. aureus,* both genes are annotated as essential^49,50^, underscoring the importance of chaperones to *S. aureus* fitness and the limitations in classifying bacterial genes into purely essential or defensive categories.

### An essential and defensive gene cotarget outperforms traditional antibiotics for eradicating established *S. aureus* biofilms

One of the major barriers to treating *S. aureus* is its ability to form recalcitrant biofilms, which can reduce antibiotic efficiency by up to a factor of 1,000^51^. To elucidate the potency of cotargeting PLPs against biofilms, established biofilms were treated with the synergistic ME+LD cotarget, a non-targeting control, or traditional antibiotics (**Fig. 5A**). The traditional antibiotics vancomycin and ampicillin demonstrated moderate biofilm removal at all concentrations, but did not increase in potency at higher doses, presumably due to the well-documented diffusion limitations in biofilms^52,53^ (**Fig. 5B**). At 15 times the minimum bactericidal concentration (MBC), vancomycin reduced biofilm abundance by 42.9±13.1% relative to the untreated control, with ampicillin achieving 34.0±16.8% reduction. Non-targeting PLPs removed 29.2±3.4% at the highest concentration, which is likely due to the activity of wild-type phages in the PLP preparation (**Fig. 5B**). In contrast to traditional antibiotics, ME+LD demonstrated a dose-dependent reduction in biofilm, and at the highest concentration, significantly outperformed both antibiotics at 15x their MBCs (**Fig. 5B**). When treated with 1 × 10^8^ transducing units (TRU), ME+LD eradicated 67.9±9.3% of the biofilm, supporting the use of cotargeting CRISPR-Cas antimicrobials as potent anti-biofilm agents.

**Figure 5:**
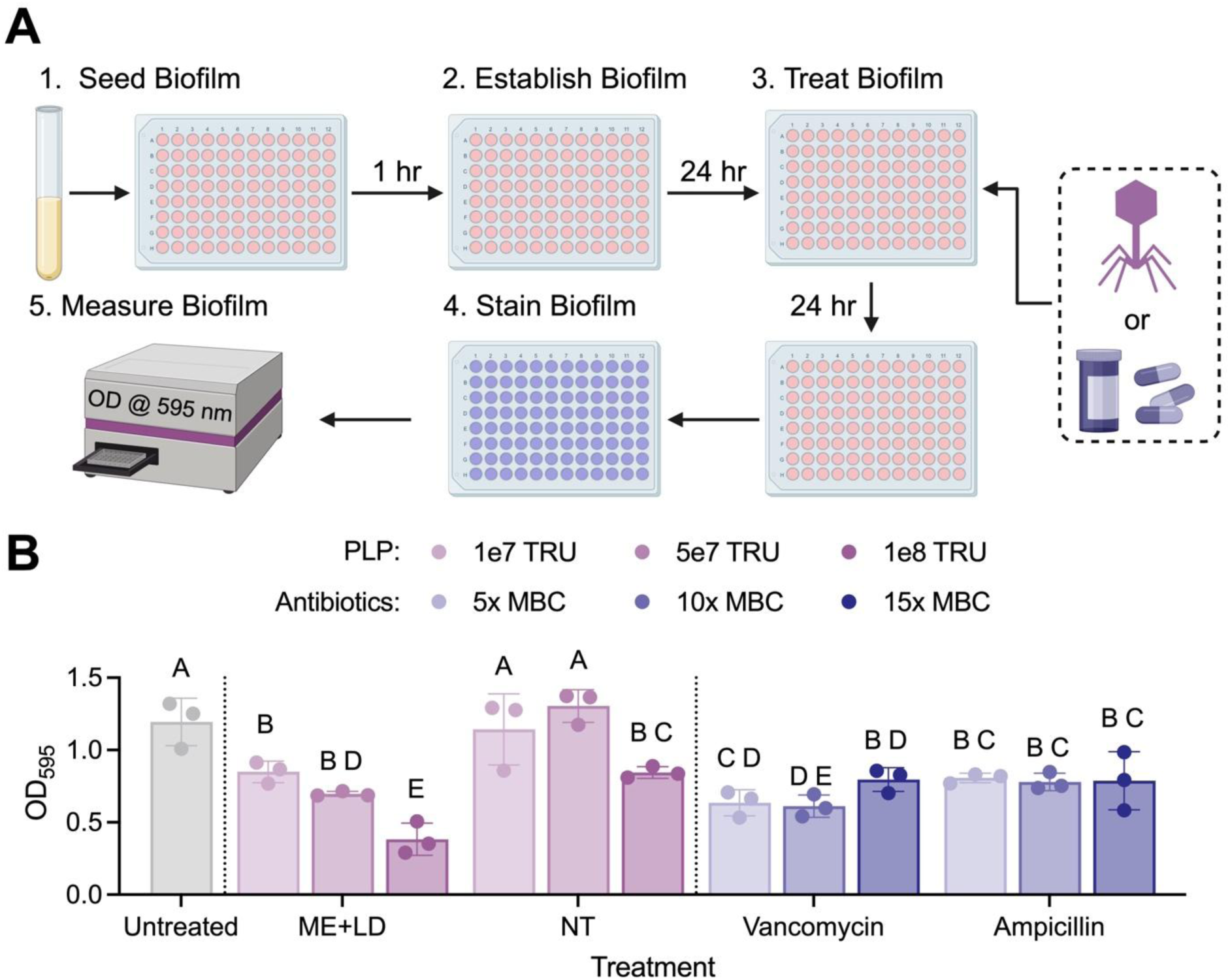
An essential and defensive gene cotarget demonstrates improved biofilm eradication over traditional antibiotics. (**A**) Experimental workflow for biofilm removal assay. (**B**) Biofilm abundances for PLP and antibiotic treatments at various concentrations. Statistical significance determined by one-way ANOVA with Fisher’s LSD test.

## CONCLUSION

This work establishes a design framework for highly potent CRISPR-Cas antimicrobials that cotarget essential and defensive genes for enhanced antimicrobial efficacy in *S. aureus*. This design strategy is made possible by the development of a novel anti-CRISPR production strain that enables the examination of cotargets across the *S. aureus* genome. Using a library depletion screen, we identified highly active guides targeting essential and defensive genes for CRISPR-Cas antimicrobial formulation in *S. aureus*. Self-targeting CRISPR antimicrobials were then generated and titrated using this anti-CRISPR production platform, which facilitates CRISPR-Cas antimicrobial development. The lethality profiles of top guides fluctuated according to targeted gene function, with proteogenesis and proteostasis emerging as particularly attractive processes for CRISPR targeting. We revealed that gene cotargeting, especially across essential and defensive pathways, significantly improves the lethality of CRISPR-Cas treatments. Defensive targets involved in signaling and stress responses were particularly effective in cotargeting strategies. In particular, simultaneous targeting of *rpoB* and *lexA* exhibited strong synergy and killing activity, despite neither gene target achieving high efficacy in isolation. When applied to biofilms, the *rpoB*-*lexA* cotarget outperformed traditional antibiotics at high concentrations, demonstrating its utility in refractory treatment scenarios. While the CRISPR screen focused on a representative set of 20 defensive and essential genes in this study, future work can expand exploration of cotargets across the genome of *S. aureus* and additional genetic backgrounds. These cotargeting strategies may also be applied to other pathogens as a generalizable framework for CRISPR-Cas antimicrobial development.

## MATERIALS AND METHODS

### Strains and plasmids

All plasmids, strains, and primers used in this study are found in Tables S1, S2, and S3, respectively. The pCasSA editing plasmid (Addgene #98211) was a gift from Quanjiang Ji and served as the backbone for the library destination vector and all phagemids^39^. The pUC19 vector was a gift from Joachim Messing (Addgene #50005) and was used as the backbone for library storage vectors^54^. The pRN111 integrative plasmid (Addgene #84463) was a gift from Reindert Nijland and served as the backbone for the AcrIIA4 integration plasmid^55^. AcrIIA4 was amplified from pCSW21, which was a gift from Joseph Bondy-Denomy (Addgene #86836)^42^.

Unless otherwise noted, all strains were grown at 37 °C with 250 rpm shaking. *E. coli* strains were cultured in lysogeny broth (LB) with 50 µg/mL kanamycin or 100 µg/mL ampicillin when appropriate. *S. aureus* strains were cultured in tryptic soy broth (TSB) or on tryptic soy agar (TSA) with 10 µg/mL chloramphenicol (Cm10) when appropriate.

### Molecular cloning

Plasmids were constructed in either *E. coli* 10-beta (NEB #C3019H) or Clean Genome LowMut (Scarab Genomics #C-6786). PCRs for cloning were performed with Phusion™ Hot Start II DNA Polymerase (Thermo Scientific™ #F549L) according to manufacturer’s recommendations. Gibson assembly reactions were performed as previously described^56^ and 1 µL was used for transformation. Ligation reactions were performed in NEB T4 DNA ligase (#M0202) and incubated at 16 °C overnight before transformation of 1 µL reaction product. Golden gate assembly reactions were performed with NEB SapI (#R0569) or BbsI-HF (#R3539) using 0.5 µL IIS enzyme and 0.5 µL T4 DNA ligase in a 10 µL reaction volume. Reactions were cycled 60 times at 37 °C and 16 °C for 5 min each, followed by 1 hr at 37 °C, 20 min at 80 °C, and finally held at 4 °C.

When annealed oligos were used for cloning, the following protocol was used: complementary oligos (1 µL each at 100 µM) were combined with 5 µL of 10x NEB T4 DNA ligase buffer (#B0202), 1 µL of NEB T4 polynucleotide kinase (#M0201), and 42 µL nuclease-free water for a final volume of 50 µL. The reaction was incubated at 37 °C for 1 hr to allow phosphorylation. After incubation, 2.5 µL of 1 M NaCl was added to the reaction, and the mixture was heated to 95 °C for 5 min to anneal oligos. Oligos were allowed to cool at room temperature for at least 2 hr. Before use, the annealed oligos were diluted to a final concentration of 0.1 µM in nuclease-free water, and 1 µL (100 fmol) was used for cloning.

#### Heat shock transformation

To heat shock transform *E. coli*, 50 µL of chemically competent cells were thawed on ice for 10 min before 1 µL of cloning reaction was added. Cells were incubated on ice for 30 min, heat shocked in a 42 °C water bath for 30 sec, then immediately returned to ice for 3 min. Next, cells were recovered in 950 µL of LB media at 37 °C for 1 hr with 250 rpm shaking. Finally, cells were pelleted by centrifugation at 8,000 xg for 3 min, resuspended in 100 µL of sterile Millipore water, plated on 1.5% LB agar with appropriate antibiotics, and incubated overnight. Transformants were confirmed by colony PCR and extracted plasmids were verified using restriction enzyme digestion and/or Sanger sequencing.

#### Library construction

The library was cloned at a depth ≥10n in *E. coli* 10-beta, where n is the library size (n = 88), using a modified CombiGEM approach^57^. To begin, the level 0 storage vector, pDDSV0, and the level 1 destination vector, pDDDV1, were cloned. pDDSV0 was cloned by digesting pUC19 with SapI and EcoRI overnight at 37 °C, followed by purification with the Omega Bio-tek E.Z.N.A Cycle Pure Kit (#D6492). Oligos DD001.f/r were annealed and ligated into the linearized pUC19 backbone, introducing a SapI stuffer cassette. To construct pDDDV1, the ts *ori* from pCasSA was replaced with a non-ts *ori* from pSGFPS1. The pCasSA backbone was amplified with DD002.f/r and the *ori* from pSGFPS1 was amplified with DD003.f/r. Fragments were assembled into pDDDV1 using Gibson assembly.

Guide RNAs were ordered as complementary forward and reverse single-stranded oligos from Eurofins Genomics containing the 20-bp gRNA sequence, a BbsI stuffer, and a unique 8-bp barcode that was generated using the nrpcalc library in a custom Python script^58^. Oligos were designed to produce an annealed product with 3-bp overhangs compatible with SapI-mediated insertion into pDDSV0. Level 0 storage vectors with added gRNA inserts were labeled level 1 storage vectors (pDDSV1). The gRNA scaffold was ordered as a dsDNA fragment with flanking BbsI cut sites and was inserted into the BbsI stuffer of pDDSV1 plasmids using Golden Gate assembly. The resultant vectors were labeled level 2 storage vectors (pDDSV2). Lastly, guide RNA cassettes were PCR amplified from pDDSV2 vectors using DD004.f/r and digested with DpnI to remove template DNA, followed by an overnight dual digest with AvrII and XhoI. Digested PCR products were cleaned with Omega Bio-tek Mag-Bind TotalPure NGS beads (#M1378) and ligated into pDDDV1 digested with SalI and XbaI.

#### Base SpCas9 phagemid cloning

To construct a base vector for CRISPR-Cas antimicrobial characterization, the *terS* gene from *ϕ*NM1 was inserted into pDDDV1. *TerS* was amplified from purified *S. aureus* SaDD0001 genomic DNA using DD005.f/r. The insert and pDDDV1 were both digested with XbaI and XhoI then ligated together to form pDD0021. Guides were added to pDD0021 using the same procedure described for pDDDV1. To generate cotargeting constructs, single-gRNA pDD0021 vectors were digested with SalI and XbaI and the second gRNA cassette was ligated in, as done for pDDDV1 and pDD0021 vectors.

#### AcrIIA4 integrative vector cloning

To create the integrative vector for inserting AcrIIA4 into *S. aureus*, the pRN111 backbone was amplified with DD006.f/r and *AcrIIA4* was amplified from pCSW21 with DD007.f/r. Both fragments were cleaned and combined using Gibson assembly to create pDD1044.

### Transformation into *S. aureus*

Electrocompetent *S. aureus* cells were prepared according to Schneewind et al.^41^. Purified plasmid DNA was added to 100 µL of electrocompetent cells and the mixture was transferred to a pre-chilled 1 mm gap electroporation cuvette before pulsing at 2,500 V, 100 Ω, 25 µF. Immediately after applying the electrical pulse, 1 mL room-temperature TSB was added directly to the cuvette, and the sample was transferred to a sterile 1.5 mL microcentrifuge tube. Samples were recovered without shaking at 30 °C for 2 hr if the plasmid contained a temperature-sensitive (ts) origin of replication (*ori*) or at 37 °C for 1 hr if it did not. Following recovery, 100 µL of cells were plated on pre-warmed 1.5% TSA plates with appropriate antibiotics and incubated overnight (37 °C) or for 36 hr (30 °C), depending on the plasmid’s origin.

Transformants were confirmed using Phire Plant Direct PCR Master Mix (Thermo Scientific™ #F160L). Briefly, a single colony was dissolved in 20 µL sterile Millipore water and 4.8 µL of the cell suspension was added to 5 µL of 2X Phire master mix and 0.1 µL of each primer (1 µM final). The reaction was thermocycled according to the manufacturer’s recommendations and the resultant amplicon was visualized on a 1% agarose gel.

### Pooled CRISPR library screen

#### Gene target selection and gRNA design

Essential and defensive gene targets were selected, and gRNAs were designed. To select candidate essential genes, all the entries for *S. aureus* in the Database of Essential Genes v15 were downloaded^37^. Essential genes that were conserved across all recorded strains of *S. aureus* were analyzed by functional categories (i.e., COGs). Essential genes were taken from each of the functional categories (10 total). Defensive genes were selected based on previous data from within the Trinh Lab and from literature, with a focus on DNA repair genes. A table of the final gene targets is provided in **Fig. 1C**. Next, guide RNA sequences were designed for each of the 20 gene targets using CASPER, with 4 guides designed for each gene^36^. The top two guides predicted by CASPERon and Azimuth 2.0 on-target scoring algorithms were chosen^59,60^. If the top two overlapped for both scores, the gRNA with the next highest CASPERon score was taken. Additionally, eight “control” guides were constructed: two guides targeting non-coding regions of the genome, two non-targeting guides (negative controls), two multitargeting guides, and two guides with established activity in *S. aureus*^39^. Plates containing the guide RNA oligos were ordered from Eurofins Genomics.

#### Library depletion assay

The finished guide library was passaged through *E. coli* IM08*β* to confer the appropriate methylation pattern to avoid restriction by *S. aureus* Newman^40^. Passaged libraries (0 hr samples) were electroporated into *S. aureus* Newman in triplicate and directly recovered into shaker flasks containing fresh selective media. Samples were taken at 6 hr and 24 hr post-transformation and plasmids were extracted, gRNA cassettes amplified with DD008.f/r, and amplicons sequenced with 2×150 paired-end Illumina reads.

#### Bioinformatics analysis

Log_2_ reductions of gRNAs were calculated using CRISPRCloud2^61^ using default settings. Counts were determined using the 20-bp gRNA sequence for both forward and reverse reads.

### AcrIIA4 integration into *S. aureus*

To create *S. aureus* SaDD0006, pDD1044 was electroporated into *S. aureus* SaDD0001 and transformants were confirmed by colony PCR. Colonies were restreaked onto TSA*_Cm_*_10_ and grown overnight at 45 °C to select for chromosomal integration of the plasmid. Large colonies were restreaked on TSA*_Cm_*_10_ and incubated another night at 45 °C. Next, colonies were cultured in TSB at 30 °C and diluted 1000x twice daily for 2.5 days (5 dilution-culture cycles), once in the morning and once in the evening. After 5 cycles, cultures were serially diluted in sterile Millipore water and 100 µL of 1×10^3^ and 1×10^4^ dilutions were plated on TSA supplemented with 100 ng/mL anhydrotetracycline (aTc100) and incubated overnight at 37 °C to select for double crossover mutants. Colonies were screened for AcrIIA4 insertion with DD009f./r and positive clones were streaked on TSA*_Cm_*_10_ and TSA to verify the loss of the plasmid. If colonies grew only in the absence of selection, overnight cultures were lysed with lysostaphin (Biosynth #ENZ-269) as previously described^41^, gDNA was purified using the Omega Bio-tek E.Z.N.A. Bacterial DNA Kit (#D3350), and the presence of AcrIIA4 was confirmed by Phusion PCR on 50 ng gDNA with DD009.f/r and Sanger sequencing with DD0010. If colonies grew on TSA*_Cm_*_10_, indicating plasmid retention, the colony was restreaked on TSA*_aTc_*_100_ and grown overnight at 45 °C. Colonies were then restreaked on selective and nonselective TSA as described above, and this process was repeated until the plasmid had been successfully cured.

### CRISPR-Cas antimicrobials

#### Generation of CRISPR phage-like particles

CRISPR phagemids were transformed into *S. aureus* SaDD0006 and confirmed by colony PCR. Confirmed colonies were grown overnight in 1 mL TSB*_Cm_*_10_. Overnight cultures were diluted 50x into 2 mL fresh selective media and grown to an OD_600_ ∼ 0.6 before being induced with 0.5 µg/mL mitomycin C. Cultures were then transferred to 30 °C with 250 rpm shaking for 16–20 hr to allow lysis to occur. To kill any residual bacteria, 1–2 drops of chloroform were added to the lysed cultures, which were then incubated at 30 °C for 15 min. Cultures were spun down at 3,000 xg for 10 min to sediment cell debris and 1.2 mL of supernatant was transferred to a sterile 1.5 mL Beckman Coulter tube (#357448) without disturbing the pellet. Lysates were spun down at 35,000 xg for 30 min at 4 °C in a Beckman Optima MAX-XP ultracentrifuge to pellet PLPs, supernatant was discarded, and the pellet was resuspended in 30 µL of SM buffer (200 mM NaCl, 10 mM MgSO_4_, 50 mM Tris-HCl, pH 7.5). Samples were stored at 4 °C until use.

#### Phagemid titer quantification

A schematic overview of this method is provided in **Fig. S2A**. A single isolated colony of *S. aureus* SaDD0006 was inoculated into 1 mL TSB supplemented with 5 mM CaCl_2_ and grown overnight at 37 °C. The overnight culture was diluted 50x and grown to an OD_600_ ∼ 1.2 in TSB with 5 mM CaCl_2_. The culture was diluted to an OD_600_ = 1.0 and 100 µL was mixed with 1 µL of PLP lysate and 99 µL of TSB with 5 mM CaCl_2_. The transduction reaction was incubated at 37 °C for 1 hr without shaking, serially diluted in sterile Millipore water, and spotted on TSA*_Cm_*_10_ plates. Plates were incubated overnight at 37 °C and TRU were enumerated the next day. TRU/mL of lysate was back calculated from TRU, the volume of lysate added, and the dilution factor from which colonies were counted. After centrifugation and resuspension, all lysates had phagemid concentrations between 10^9^ and 10^11^ TRU/mL, with a median concentration of 7.6×10^9^ TRU/mL (**Fig. S2B**).

#### Transduction killing assay

A schematic overview of this method is provided in **Fig. 3A**. A single isolated colony of *S. aureus* SaDD0001 was inoculated into 1 mL TSB supplemented with 5 mM CaCl_2_ and grown overnight at 37 °C. The overnight culture was diluted 50x and grown to an OD_600_ ∼ 0.6 in TSB with 5 mM CaCl_2_. The culture was diluted to an OD_600_ = 0.04 and 100 µL was combined with PLP lysate at an MOI = 1 using TRU/mL values and the assumption that 1 OD_600_ = 1 × 10^8^ cells/mL^41^. Lysate volume was adjusted to 100 µL in TSB with 5 mM CaCl_2_ and the transduction reaction was incubated at 37 °C for 1 hr without shaking and then 180 µL was plated on TSA*_Cm_*_10_ plates. Plates were incubated at 37 °C overnight and surviving colonies were enumerated the next day. TRU/mL was calculated as previously described and log_10_ reduction was calculated relative to the non-targeting control. For antibiotic controls, vancomycin and chloramphenicol were added to cultures at twice their minimum inhibitory concentrations^62^. Cells were treated for 4 hr at 37 °C, then plated on TSA and grown overnight to enumerate surviving colonies. Log_10_ reduction was calculated relative to an untreated control.

#### Interaction score analysis

Interaction score analysis was performed using a modified gene cotargeting interaction formula^29^. The expected log_10_ reduction (LR) of a cotarget with guides i and j was assumed to be equal to the average of the LR values for the two self constructs for guides i and j.

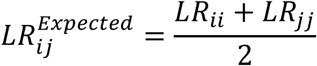

The interaction score (IS) was then calculated by subtracting the expected LR for a cotarget from the observed LR of the cotarget.

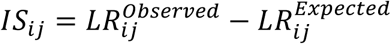

Under this formulation, cotargets with interaction scores greater than zero are considered synergistic, while cotargets with interaction scores less than zero are considered antagonistic.

#### Biofilm killing assay

*S. aureus* Newman was grown overnight and diluted 50x into fresh media the following day. Cultures were grown to mid-exponential phase (OD ∼ 0.6) and diluted back to exactly 1 × 10^6^ cells/mL in RPMI 1640 media (Gibco #31800-022) supplemented with 25 mM HEPES and 2 g/L sodium bicarbonate (hereafter, simply “RPMI”) that was pre-warmed at 37 °C. 100 µL of cells were aliquoted into each well of a sterile, surface-treated 96-well plate (VWR #10062-900) for a total of 1 × 10^5^ cells per well. Plates were incubated statically for 1 hr at 37 °C to allow cells to adhere to the microplate. Supernatant was then removed and wells were gently washed twice with 200 µL sterile phosphate-buffered saline (PBS) to remove non-adherent cells. Next, 200 µL of fresh, pre-warmed RPMI was added to each well. Microplates were statically incubated at 37 °C for 24 hr to establish biofilms. After 24 hr, supernatant was again removed and wells were gently washed twice with 200 µL sterile PBS. PLP and antibiotic treatments were diluted to the appropriate concentrations in RPMI and added to the established biofilms. RPMI alone was added to the untreated controls. Biofilms were treated for 24 hr at 37 °C, then supernatant removal and PBS washes were repeated once more. Plates were air dried at 37 °C for 20–30 min before biofilms were stained with 1% crystal violet. Biofilms were stained for 20 min at room temperature, crystal violet was removed, and biofilms were washed four times with Millipore water. Plates were again air dried at 37 °C for 20–30 min. Finally, biofilms were destained by incubating with 33% acetic acid for 45 min. 150 µL of destained crystal violet was transferred to a new microplate and the absorbance was measured at 595 nm to determine the biofilm abundance. Readings were normalized by subtracting the average values of no-cell controls.

## ACKNOWLEDGEMENTS

This research was funded by the DARPA YFA award and Director Fellowship (D17AP00023). The authors would like to sincerely thank Dr. Ed Wright and the Bioanalytical Resource Facility at the University of Tennessee for the use of the Beckman Optima MAX-XP ultracentrifuge. The following reagents were obtained through BEI Resources, NIAID, NIH: *Staphylococcus aureus* Fluorescent Reporter Plasmid pSGFPS1, Recombinant in *Staphylococcus aureus*, NR-51163. The mention of trade names or commercial products in this article is solely for the purpose of providing scientific information and does not imply recommendation or endorsement by the funding agency.

