## Supplementary Figure S1-S3 and Table S1-S3 for "Synergistic CRISPR-Cas Antimicrobials through Essential and Defensive Gene Cotargeting in *Staphylococcus aureus*"

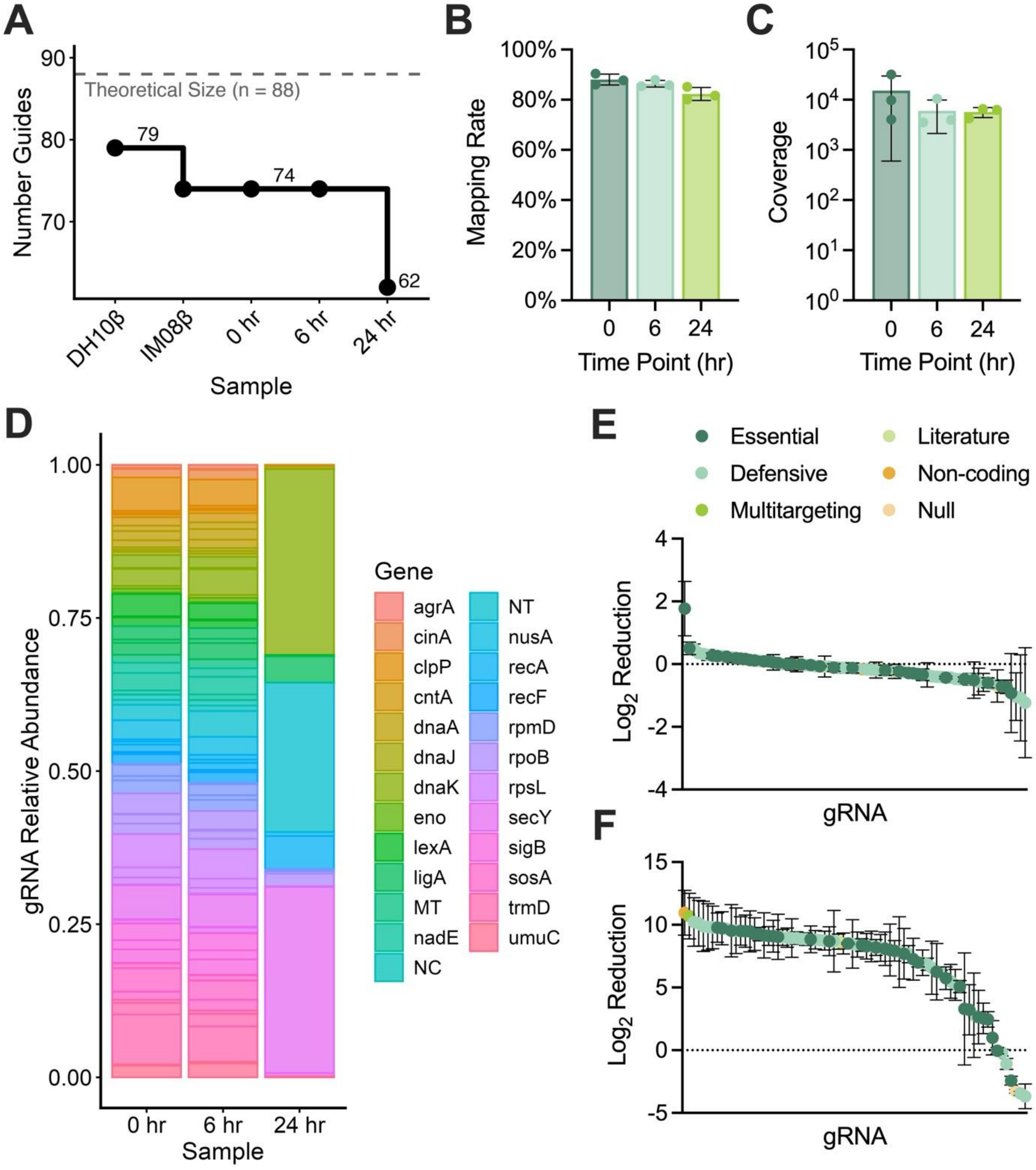


**Figure S1:** Quality statistics of pooled CRISPR library screen. (**A**) Number of guides detected in the library for each sample. (**B**) Mapping rate for each time point. The mean of each time point was ≥80%. (**C**) Library coverage by sequencing reads at each time point. (**D**) Relative abundance of gRNAs at 0, 6, and 24 hr time points. (**E–F**) $Log_2$ reductions of each gRNA relative to the 0 hr time point at (**E**) 6 hr and (**F**) 24 hr.

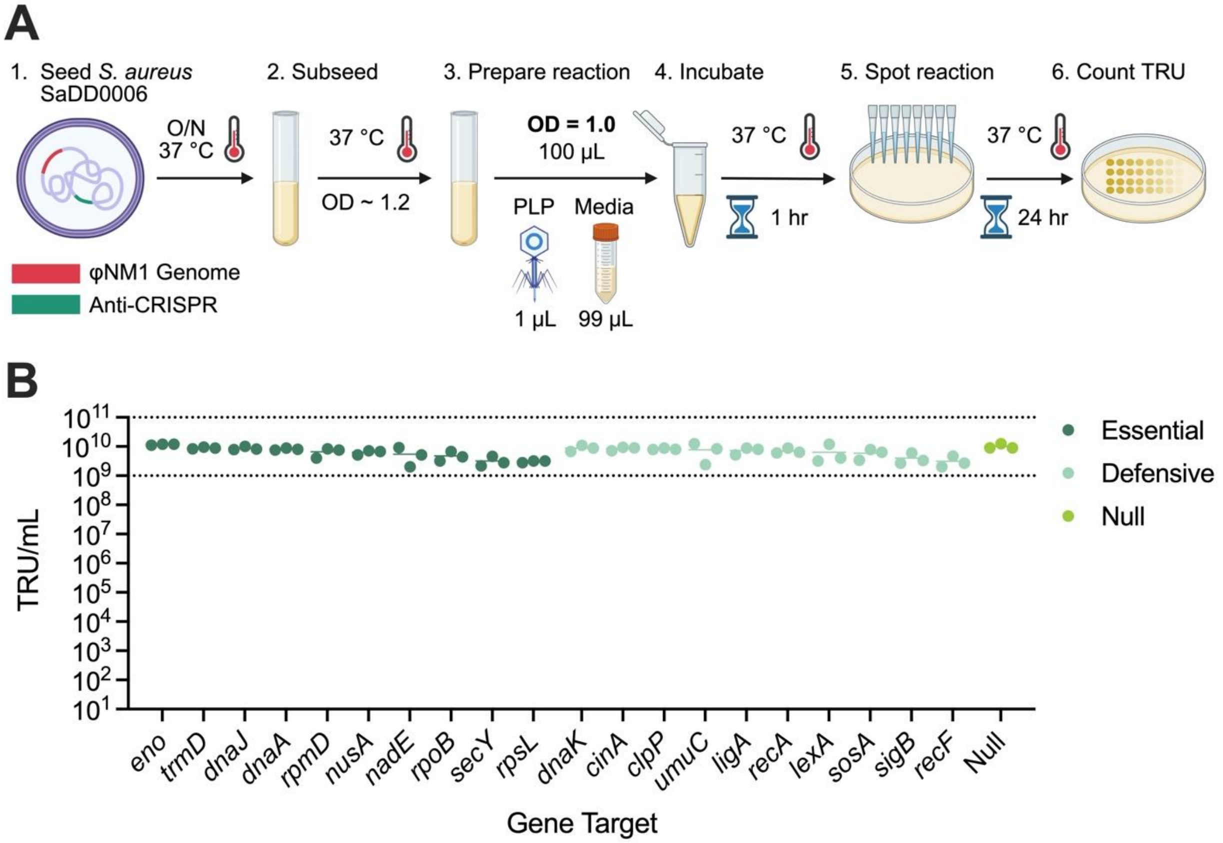


**Figure S2:** *S. aureus* SaDD0006 enables accurate, high-throughput quantification of CRISPR phagemids in phage-like particles (PLPs). (**A**) Workflow for quantifying phagemid concentration. (**B**) Phagemid concentration of PLPs harboring single gRNAs.

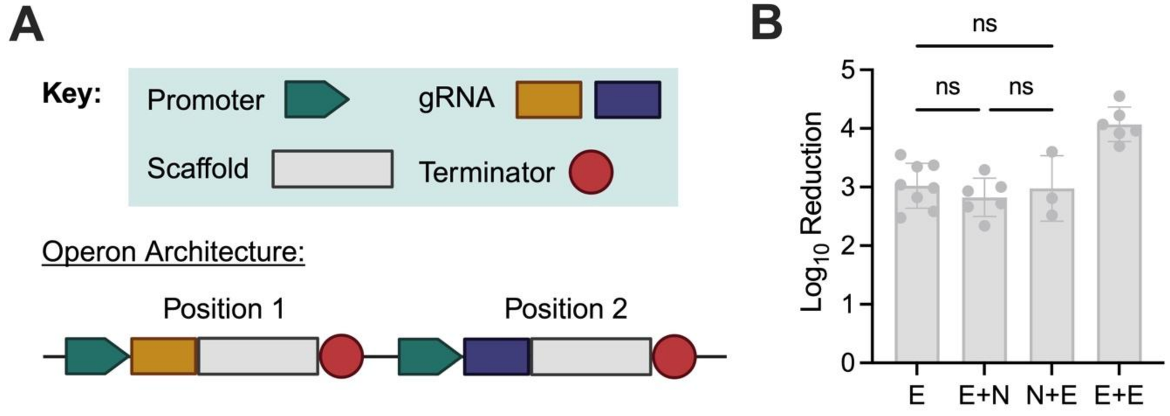


**Figure S3:** Discrete transcriptional units mitigate position-dependent effects in multiplexed constructs. (**A**) Schematic of multiplexed gRNA operon architecture. (**B**) $Log_{10}$ reduction of an essential gRNA alone (E), in position 1 with a null gRNA (E+N), in position 2 with a null gRNA (N+E), or in both positions 1 and 2 (E+E). Statistical significance determined by one-way ANOVA.

**Table S1:** List of plasmids used in this study.

| Plasmid ID | Parent Plasmid | Role in Study | Gene Targets | gRNA Classification | Sources |
|---|---|---|---|---|---|
| pCasSA | | Base plasmid | | | Addgene #98211 |
| pRN111 | | Base plasmid | | | Addgene #84463 |
| pUC19 | | Base plasmid | | | Addgene #50005 |
| pCSW21 | | AcrIIA4 source | | | Addgene #86836 |
| pSGFPS1 | | Non-ts *S. aureus ori* source | | | BEI (NR-51163) |
| pDD1044 | pRN111 | AcrIIA4 integrative vector | | | This Study |
| pDDSV0 | pUC19 | Level 0 gRNA storage vector | | | This Study |
| pDDSV1 | pDDSV0 | Level 1 gRNA storage vector | | | This Study |
| pDDSV2 | pDDSV1 | Level 2 gRNA storage vector | | | This Study |
| pDDDV1 | pCasSA | Level 1 gRNA destination vector | | | This Study |
| pDD0021 | pDDDV1 | Base SpCas9 phagemid | | | This Study |
| pDDSV1-trmD-4 | pDDSV0 | Level 1 gRNA storage vector | *trmD* | E | This Study |
| pDDSV1-rpsL-4 | pDDSV0 | Level 1 gRNA storage vector | *rpsL* | E | This Study |
| pDDSV1-dnaJ-1 | pDDSV0 | Level 1 gRNA storage vector | *dnaJ* | E | This Study |
| pDDSV1-rpoB-3 | pDDSV0 | Level 1 gRNA storage vector | *rpoB* | E | This Study |
| pDDSV1-secY-2 | pDDSV0 | Level 1 gRNA storage vector | *secY* | E | This Study |
| pDDSV1-eno-4 | pDDSV0 | Level 1 gRNA storage vector | *eno* | E | This Study |
| pDDSV1-nadE-1 | pDDSV0 | Level 1 gRNA storage vector | *nadE* | E | This Study |
| pDDSV1-rpmD-1 | pDDSV0 | Level 1 gRNA storage vector | *rpmD* | E | This Study |
| pDDSV1-dnaA-1 | pDDSV0 | Level 1 gRNA storage vector | *dnaA* | E | This Study |
| pDDSV1-nusA-3 | pDDSV0 | Level 1 gRNA storage vector | *nusA* | E | This Study |
| pDDSV1-recA-2 | pDDSV0 | Level 1 gRNA storage vector | *recA* | D | This Study |
| pDDSV1-clpP-4 | pDDSV0 | Level 1 gRNA storage vector | *clpP* | D | This Study |
| pDDSV1-dnaK-1 | pDDSV0 | Level 1 gRNA storage vector | *dnaK* | D | This Study |
| pDDSV1-lexA-1 | pDDSV0 | Level 1 gRNA storage vector | *lexA* | D | This Study |

| | | | | | |
|---|---|---|---|---|---|
| pDDSV1-sigB-4 | pDDSV0 | Level 1 gRNA storage vector | *sigB* | D | This Study |
| pDDSV1-sosA-1 | pDDSV0 | Level 1 gRNA storage vector | *sosA* | D | This Study |
| pDDSV1-ligA-1 | pDDSV0 | Level 1 gRNA storage vector | *ligA* | D | This Study |
| pDDSV1-recF-4 | pDDSV0 | Level 1 gRNA storage vector | *recF* | D | This Study |
| pDDSV1-umuC-2 | pDDSV0 | Level 1 gRNA storage vector | *umuC* | D | This Study |
| pDDSV1-cinA-4 | pDDSV0 | Level 1 gRNA storage vector | *cinA* | D | This Study |
| pDDSV1-NT | pDDSV0 | Level 1 gRNA storage vector | non-targeting (NT) | Null | This Study |
| pDDSV2-trmD-4 | pDDSV1 | Level 2 gRNA storage vector | *trmD* | E | This Study |
| pDDSV2-rpsL-4 | pDDSV1 | Level 2 gRNA storage vector | *rpsL* | E | This Study |
| pDDSV2-dnaJ-1 | pDDSV1 | Level 2 gRNA storage vector | *dnaJ* | E | This Study |
| pDDSV2-rpoB-3 | pDDSV1 | Level 2 gRNA storage vector | *rpoB* | E | This Study |
| pDDSV2-secY-2 | pDDSV1 | Level 2 gRNA storage vector | *secY* | E | This Study |
| pDDSV2-eno-4 | pDDSV1 | Level 2 gRNA storage vector | *eno* | E | This Study |
| pDDSV2-nadE-1 | pDDSV1 | Level 2 gRNA storage vector | *nadE* | E | This Study |
| pDDSV2-rpmD-1 | pDDSV1 | Level 2 gRNA storage vector | *rpmD* | E | This Study |
| pDDSV2-dnaA-1 | pDDSV1 | Level 2 gRNA storage vector | *dnaA* | E | This Study |
| pDDSV2-nusA-3 | pDDSV1 | Level 2 gRNA storage vector | *nusA* | E | This Study |
| pDDSV2-recA-2 | pDDSV1 | Level 2 gRNA storage vector | *recA* | D | This Study |
| pDDSV2-clpP-4 | pDDSV1 | Level 2 gRNA storage vector | *clpP* | D | This Study |
| pDDSV2-dnaK-1 | pDDSV1 | Level 2 gRNA storage vector | *dnaK* | D | This Study |
| pDDSV2-lexA-1 | pDDSV1 | Level 2 gRNA storage vector | *lexA* | D | This Study |
| pDDSV2-sigB-4 | pDDSV1 | Level 2 gRNA storage vector | *sigB* | D | This Study |
| pDDSV2-sosA-1 | pDDSV1 | Level 2 gRNA storage vector | *sosA* | D | This Study |
| pDDSV2-ligA-1 | pDDSV1 | Level 2 gRNA storage vector | *ligA* | D | This Study |
| pDDSV2-recF-4 | pDDSV1 | Level 2 gRNA storage vector | *recF* | D | This Study |
| pDDSV2-umuC-2 | pDDSV1 | Level 2 gRNA storage vector | *umuC* | D | This Study |
| pDDSV2-cinA-4 | pDDSV1 | Level 2 gRNA storage vector | *cinA* | D | This Study |
| pDDSV2-NT | pDDSV1 | Level 2 gRNA storage vector | non-targeting (NT) | Null | This Study |
| pDD0021-trmD-2 | pDD0021 | Monoplexed phagemid | *trmD* | E | This Study |

| | | | | | |
|---|---|---|---|---|---|
| pDD0021-trmD-4 | pDD0021 | Monoplexed phagemid | *trmD* | E | This Study |
| pDD0021-rpsL-4 | pDD0021 | Monoplexed phagemid | *rpsL* | E | This Study |
| pDD0021-dnaJ-1 | pDD0021 | Monoplexed phagemid | *dnaJ* | E | This Study |
| pDD0021-rpoB-3 | pDD0021 | Monoplexed phagemid | *rpoB* | E | This Study |
| pDD0021-secY-2 | pDD0021 | Monoplexed phagemid | *secY* | E | This Study |
| pDD0021-eno-4 | pDD0021 | Monoplexed phagemid | *eno* | E | This Study |
| pDD0021-nadE-1 | pDD0021 | Monoplexed phagemid | *nadE* | E | This Study |
| pDD0021-rpmD-1 | pDD0021 | Monoplexed phagemid | *rpmD* | E | This Study |
| pDD0021-dnaA-1 | pDD0021 | Monoplexed phagemid | *dnaA* | E | This Study |
| pDD0021-nusA-3 | pDD0021 | Monoplexed phagemid | *nusA* | E | This Study |
| pDD0021-recA-2 | pDD0021 | Monoplexed phagemid | *recA* | D | This Study |
| pDD0021-clpP-4 | pDD0021 | Monoplexed phagemid | *clpP* | D | This Study |
| pDD0021-dnaK-1 | pDD0021 | Monoplexed phagemid | *dnaK* | D | This Study |
| pDD0021-lexA-1 | pDD0021 | Monoplexed phagemid | *lexA* | D | This Study |
| pDD0021-sigB-4 | pDD0021 | Monoplexed phagemid | *sigB* | D | This Study |
| pDD0021-sosA-1 | pDD0021 | Monoplexed phagemid | *sosA* | D | This Study |
| pDD0021-ligA-1 | pDD0021 | Monoplexed phagemid | *ligA* | D | This Study |
| pDD0021-recF-4 | pDD0021 | Monoplexed phagemid | *recF* | D | This Study |
| pDD0021-umuC-2 | pDD0021 | Monoplexed phagemid | *umuC* | D | This Study |
| pDD0021-cinA-4 | pDD0021 | Monoplexed phagemid | *cinA* | D | This Study |
| pDD0021-NT | pDD0021 | Monoplexed phagemid | non-targeting (NT) | Null | This Study |
| pDD0021-HE+HD | pDD0021-secY-2 | Multiplexed phagemid | *secY* and *clpP* | E+D | This Study |
| pDD0021-HE+ME | pDD0021-secY-2 | Multiplexed phagemid | *secY* and *rpoB* | E+E | This Study |
| pDD0021-HE+MD | pDD0021-secY-2 | Multiplexed phagemid | *secY* and *dnaK* | E+D | This Study |
| pDD0021-HE+LE | pDD0021-secY-2 | Multiplexed phagemid | *secY* and *trmD* | E+E | This Study |
| pDD0021-HE+LD | pDD0021-secY-2 | Multiplexed phagemid | *secY* and *lexA* | E+D | This Study |
| pDD0021-HD+ME | pDD0021-clpP-4 | Multiplexed phagemid | *clpP* and *rpoB* | E+D | This Study |
| pDD0021-HD+MD | pDD0021-clpP-4 | Multiplexed phagemid | *clpP* and *dnaK* | D+D | This Study |
| pDD0021-HD+LE | pDD0021-clpP-4 | Multiplexed phagemid | *clpP* and *trmD* | E+D | This Study |

| | | | | | |
|---|---|---|---|---|---|
| pDD0021-HD+LD | pDD0021-clpP-4 | Multiplexed phagemid | *clpP* and *lexA* | D+D | This Study |
| pDD0021-ME+MD | pDD0021-rpoB-3 | Multiplexed phagemid | *rpoB* and *dnaK* | E+D | This Study |
| pDD0021-ME+LE | pDD0021-rpoB-3 | Multiplexed phagemid | *rpoB* and *trmD* | E+E | This Study |
| pDD0021-ME+LD | pDD0021-rpoB-3 | Multiplexed phagemid | *rpoB* and *lexA* | E+D | This Study |
| pDD0021-MD+LE | pDD0021-dnaK-1 | Multiplexed phagemid | *dnaK* and *trmD* | E+D | This Study |
| pDD0021-MD+LD | pDD0021-dnaK-1 | Multiplexed phagemid | *dnaK* and *lexA* | D+D | This Study |
| pDD0021-LE+LD | pDD0021-trmD-4 | Multiplexed phagemid | *trmD* and *lexA* | E+D | This Study |
| pDD0021-HE+HE | pDD0021-secY-2 | Multiplexed phagemid | *secY* and *secY* | Self | This Study |
| pDD0021-HD+HD | pDD0021-clpP-4 | Multiplexed phagemid | *clpP* and *clpP* | Self | This Study |
| pDD0021-ME+ME | pDD0021-rpoB-3 | Multiplexed phagemid | *rpoB* and *rpoB* | Self | This Study |
| pDD0021-MD+MD | pDD0021-dnaK-1 | Multiplexed phagemid | *dnaK* and *dnaK* | Self | This Study |
| pDD0021-LE+LE | pDD0021-trmD-4 | Multiplexed phagemid | *trmD* and *trmD* | Self | This Study |
| pDD0021-LD+LD | pDD0021-lexA-1 | Multiplexed phagemid | *lexA* and *lexA* | Self | This Study |
| pDD0021-NT+NT | pDD0021-NT | Multiplexed phagemid | NT and NT | Null | This Study |
| pDD0021-trmD-2+NT | pDD0021-trmD-2 | Control for gRNA position effects | *trmD* and NT | E+N | This Study |
| pDD0021-NT+trmD-2 | pDD0021-NT | Control for gRNA position effects | NT and *trmD* | N+E | This Study |
| pDD0021-trmD-2+trmD-2 | pDD0021-trmD-2 | Control for gRNA position effects | *trmD* and *trmD* | Self/(E+E, Fig. S3B) | This Study |

**Table S2:** List of strains used in this study

| Strain | Parent Strain | Relevant Features | Source |
|---|---|---|---|
| *E. coli* 10-beta | *E. coli* DH10β | | NEB (C3019H) |
| *E. coli* IM08β | *E. coli* DH10β | Δ*dcm*, SA08BΩPN25-*hsdS* (CC1-1) of MW2 integrated between *essQ*/*cspB* | Monk et al. 2015 |
| E. coli Clean Genome LowMut | | IS element-deficient | Scarab Genomics (C-6786) |
| *S. aureus* RN4220 | | prophage-deficient, restriction-deficient, *agrA* mutant | BEI (NR-45946) |
| *S. aureus* Newman | | | ATCC (25904) |
| *S. aureus* SaDD0001 | *S. aureus* RN4220 | single $\phi$NM1 lysogen | This Study |
| *S. aureus* SaDD0006 | *S. aureus* SaDD0001 | single $\phi$NM1 lysogen, $P_{SarAP1}$-*AcrIIA4*-$T_{lambda}$ inserted between NWMN_0029/0030 | This Study |

**Table S3:** List of primers used in this study

| Name | Sequence | Purpose |
|---|---|---|
| DD_138.f | ATTTCGAAAAGCAAAATTATTATTTAAGACA TTAGCTTTGTTATCTTGGTTACCCG | Forward for amplifying pCasSA BB for *ori* swap |
| DD_138.r | TGATAATCATATTTTTCTTCCATCTCGTCAGG TATTTACCACAACAGTACGC | Reverse for amplifying pCasSA BB for *ori* swap |
| DD_139.f | TGACGAGATGGAAGAAAAATATG | Forward for amplifying pSGFPS1's *S. aureus* rep machinery |
| DD_139.r | CTAATGTCTTAAATAATAATTTTGCTTTTCG | Reverse for amplifying pSGFPS1's *S. aureus* rep machinery |
| DD_150.f | AGCCTCGAGAGAGTTTGCAAAATATACAGG GGATTATATATAATGGAAAGGAAGAGCGAG CTCTTCTCCTAGG | Forward oligo for making SV0 |
| DD_150.r | AATTCCTAGGAGAAGAGCTCGCTCTTCCTTT CCATTATATATAATCCCCTGTATATTTTGCA AACTCTCTCGAG | Reverse oligo for making SV0 |
| DD_172.f | CTAGTCTAGAGCGAGAGTTGGTGGTAAATG | Forward for *terS* gene, XbaI overhang |
| DD_172.r | CCGCTCGAGTTAACTTTCGTCATCGTACTCA CC | Reverse for *terS* gene, XhoI overhang |
| DD_194.f | ACACTCTTTCCCTACACGACGCTCTTCCGAT CTCCTGCGTTGTCGAGAGAG | Forward for amplifying sequencing amplicon from pDDDV1 libraries |
| DD_194.r | GACTGGAGTTCAGACGTGTGCTCTTCCGATC TTTCTGACTCGAGCATTCTAGG | Reverse for amplifying sequencing amplicon from pDDDV1 libraries |
| DD_218.f | GGATGTCCTGTAGCAAATGTTG | Forward for amplifying insertion locus for integrative anti-CRISPR vector |
| DD_218.r | CGCCAAAAGATGCTTTATTTGAC | Reverse for amplifying insertion locus for integrative anti-CRISPR vector |
| DD_219 | TATGTCACTTATCCTTTTGGAAATG | Sequencing primer for insertion locus for integrative anti-CRISPR vector |
| DD_231.f | AAAAAGTGAGTTGAACTAAGAATTCGTAAT CATGTCATAGCTGTTT | Forward for amplifying pRN111 backbone for AcrIIA4 insertion |
| DD_231.r | CTCTAATTAAGTCATTAATATTCATATGTAT ACATCCTCCTAAGGTACCCG | Reverse for amplifying pRN111 backbone for AcrIIA4 insertion |

| | | |
|---|---|---|
| DD_232.f | GGTACCTTAGGAGGATGTATACATATGAATATTAATGACTTAATTAGAGAAATCAAAAAC | Forward for amplifying AcrIIA4 for Gibson insertion into pRN111 |
| DD_232.r | ATGACATGATTACGAATTCTTAGTTCAACTCACTTTTTAAGGTGATT | Reverse for amplifying AcrIIA4 for Gibson insertion into pRN111 |
| DD_233.f | AGCGAGTCAGTGAGCGAG | Forward for amplifying gRNA cassette from SV2 vector |
| DD_233.r | TGTAAAACGACGGCCAGT | Reverse for amplifying gRNA cassette from SV2 vector |
| cinA-1F | AAAACCGAAAGAAATGCAACCAAGTTTGGGTCTTCGAGAAGACCTATTCATGATGGT | Forward for cinA-1 gRNA |
| cinA-1R | AGGACCATCATGAATAGGTCTTCTCGAAGACCCAAACTTGGTTGCATTTCTTTCGGT | Reverse for cinA-1 gRNA |
| cinA-2F | AAAACTTCATGACTTCCCGCCAAGTTTGGGTCTTCGAGAAGACCTATTCCGCATTTT | Forward for cinA-2 gRNA |
| cinA-2R | AGGAAAATGCGGAATAGGTCTTCTCGAAGACCCAAACTTGGCGGGAAGTCATGAAGT | Reverse for cinA-2 gRNA |
| cinA-3F | AAAGGTCAAATCGCTAATACCAAGTTTGGGTCTTCGAGAAGACCTATTCAGGTCAAT | Forward for cinA-3 gRNA |
| cinA-3R | AGGATTGACCTGAATAGGTCTTCTCGAAGACCCAAACTTGGTATTAGCGATTTGACC | Reverse for cinA-3 gRNA |
| cinA-4F | AAATGACTTAACGAAGCATACAGGTTTGGGTCTTCGAGAAGACCTATTCTTACCGTT | Forward for cinA-4 gRNA |
| cinA-4R | AGGAACGGTAAGAATAGGTCTTCTCGAAGACCCAAACCTGTATGCTTCGTTAAGTCA | Reverse for cinA-4 gRNA |
| clpP-1F | AAAATCGGTATGGCTGCATCAATGTTTGGGTCTTCGAGAAGACCTATTCGCTGGCAA | Forward for clpP-1 gRNA |
| clpP-1R | AGGTTGCCAGCGAATAGGTCTTCTCGAAGACCCAAACATTGATGCAGCCATACCGAT | Reverse for clpP-1 gRNA |
| clpP-2F | AAACAGTACGCTCTGATAAAATGGTTTGGGTCTTCGAGAAGACCTATTCTGTAAGCG | Forward for clpP-2 gRNA |
| clpP-2R | AGGCGCTTACAGAATAGGTCTTCTCGAAGACCCAAACCATTTTATCAGAGCGTACTG | Reverse for clpP-2 gRNA |
| clpP-3F | AAAGTTATTGAAACAACAAACCGGTTTGGGTCTTCGAGAAGACCTATTCCCTGATTC | Forward for clpP-3 gRNA |

| | | |
|---|---|---|
| clpP-3R | AGGGAATCAGGGAATAGGTCTTCTCGAAGA CCCAAACCGGTTTGTTGTTTCAATAAC | Reverse for clpP-3 gRNA |
| clpP-4F | AAATATCATATGCACGTTCACCGGTTTGGGT CTTCGAGAAGACCTATTCTTATCCCC | Forward for clpP-4 gRNA |
| clpP-4R | AGGGGGGATAAGAATAGGTCTTCTCGAAGA CCCAAACCGGTGAACGTGCATATGATA | Reverse for clpP-4 gRNA |
| dnaA-1F | AAAATCTATGGAGGTGTTGGTTTGTTTGGGT CTTCGAGAAGACCTATTCAGTCTGAA | Forward for dnaA-1 gRNA |
| dnaA-1R | AGGTTCAGACTGAATAGGTCTTCTCGAAGAC CCAAACAAACCAACACCTCCATAGAT | Reverse for dnaA-1 gRNA |
| dnaA-2F | AAAAACTCGCTGCATGTGGAAAGGTTTGGG TCTTCGAGAAGACCTATTCTGGACAAT | Forward for dnaA-2 gRNA |
| dnaA-2R | AGGATTGTCCAGAATAGGTCTTCTCGAAGAC CCAAACCTTTCCACATGCAGCGAGTT | Reverse for dnaA-2 gRNA |
| dnaA-3F | AAAAAGTACATAGCTATTTGACGGTTTGGGT CTTCGAGAAGACCTATTCTACGATCA | Forward for dnaA-3 gRNA |
| dnaA-3R | AGGTGATCGTAGAATAGGTCTTCTCGAAGAC CCAAACCGTCAAATAGCTATGTACTT | Reverse for dnaA-3 gRNA |
| dnaA-4F | AAAACCTTCTACTGAAACAACTGGTTTGGGT CTTCGAGAAGACCTATTCTGGTGGTC | Forward for dnaA-4 gRNA |
| dnaA-4R | AGGGACCACCAGAATAGGTCTTCTCGAAGA CCCAAACCAGTTGTTTCAGTAGAAGGT | Reverse for dnaA-4 gRNA |
| dnaJ-1F | AAATTTGGTGGACAAGGATTCAAGTTTGGGT CTTCGAGAAGACCTATTCCCTGGGTA | Forward for dnaJ-1 gRNA |
| dnaJ-1R | AGGTACCCAGGGAATAGGTCTTCTCGAAGA CCCAAACTTGAATCCTTGTCCACCAAA | Reverse for dnaJ-1 gRNA |
| dnaJ-2F | AAATTTGGCGGTTTTAGTGGCTTGTTTGGGT CTTCGAGAAGACCTATTCAATCTGAC | Forward for dnaJ-2 gRNA |
| dnaJ-2R | AGGGTCAGATTGAATAGGTCTTCTCGAAGAC CCAAACAAGCCACTAAAACCGCCAAA | Reverse for dnaJ-2 gRNA |
| dnaJ-4F | AAAGTAACATGCGAAACATGTCAGTTTGGG TCTTCGAGAAGACCTATTCGCCCATTC | Forward for dnaJ-4 gRNA |

| | | |
|---|---|---|
| dnaJ-4R | AGGGAATGGGCGAATAGGTCTTCTCGAAGA CCCAAACTGACATGTTTCGCATGTTAC | Reverse for dnaJ-4 gRNA |
| dnaK-1F | AAAATGGGAGCTGCAATCCAAGGGTTTGGG TCTTCGAGAAGACCTATTCGCTTGGCA | Forward for dnaK-1 gRNA |
| dnaK-1R | AGGTGCCAAGCGAATAGGTCTTCTCGAAGA CCCAAACCCTTGGATTGCAGCTCCCAT | Reverse for dnaK-1 gRNA |
| dnaK-2F | AAACTTGTTTTTGACTTAGGTGGGTTTGGGT CTTCGAGAAGACCTATTCGCGATTCC | Forward for dnaK-2 gRNA |
| dnaK-2R | AGGGGAATCGCGAATAGGTCTTCTCGAAGA CCCAAACCCACCTAAGTCAAAAACAAG | Reverse for dnaK-2 gRNA |
| dnaK-3F | AAAACCGCCAAGTTTGTTGTCACGTTTGGGT CTTCGAGAAGACCTATTCCTTTGAGT | Forward for dnaK-3 gRNA |
| dnaK-3R | AGGACTCAAAGGAATAGGTCTTCTCGAAGA CCCAAACGTGACAACAAACTTGGCGGT | Reverse for dnaK-3 gRNA |
| dnaK-4F | AAAGATATTCCACCAGCTGAACGGTTTGGGT CTTCGAGAAGACCTATTCCGTAAGTA | Forward for dnaK-4 gRNA |
| dnaK-4R | AGGTACTTACGGAATAGGTCTTCTCGAAGAC CCAAACCGTTCAGCTGGTGGAATATC | Reverse for dnaK-4 gRNA |
| eno-1F | AAAATGGACGAAAACGACTGGGAGTTTGGG TCTTCGAGAAGACCTATTCCCCTTGTC | Forward for eno-1 gRNA |
| eno-1R | AGGGACAAGGGGAATAGGTCTTCTCGAAGA CCCAAACTCCCAGTCGTTTTCGTCCAT | Reverse for eno-1 gRNA |
| eno-2F | AAAACTGCAGTAGGTGACGAAGGGTTTGGG TCTTCGAGAAGACCTATTCATAGCCTT | Forward for eno-2 gRNA |
| eno-2R | AGGAAGGCTATGAATAGGTCTTCTCGAAGA CCCAAACCCTTCGTCACCTACTGCAGT | Reverse for eno-2 gRNA |
| eno-3F | AAAACGAAAACGACTGGGATGGTGTTTGGG TCTTCGAGAAGACCTATTCACATAGCA | Forward for eno-3 gRNA |
| eno-3R | AGGTGCTATGTGAATAGGTCTTCTCGAAGAC CCAAACACCATCCCAGTCGTTTTCGT | Reverse for eno-3 gRNA |
| eno-4F | AAACGTGGTTTAGAAACTGCAGTGTTTGGGT CTTCGAGAAGACCTATTCTTTCTGCT | Forward for eno-4 gRNA |

| | | |
|---|---|---|
| eno-4R | AGGAGCAGAAAGAATAGGTCTTCTCGAAGACCCAAACACTGCAGTTTCTAAACCACG | Reverse for eno-4 gRNA |
| lexA-1F | AAACCGCCTAGTGTTCGCGAAATGTTTGGGTCTTCGAGAAGACCTATTCCGATACAA | Forward for lexA-1 gRNA |
| lexA-1R | AGGTTGTATCGGAATAGGTCTTCTCGAAGACCCAAACATTTCGCGAACACTAGGCGG | Reverse for lexA-1 gRNA |
| lexA-2F | AAAATTGAGGCTGGTATATTAGAGTTTGGGTCTTCGAGAAGACCTATTCGGAGGCAT | Forward for lexA-2 gRNA |
| lexA-2R | AGGATGCCTCCGAATAGGTCTTCTCGAAGACCCAAACTCTAATATACCAGCCTCAAT | Reverse for lexA-2 gRNA |
| lexA-3F | AAACGTAGGCGACAGTATGATTGGTTTGGGTCTTCGAGAAGACCTATTCTCTGCGAA | Forward for lexA-3 gRNA |
| lexA-3R | AGGTTCGCAGAGAATAGGTCTTCTCGAAGACCCAAACCAATCATACTGTCGCCTACG | Reverse for lexA-3 gRNA |
| lexA-4F | AAACCAATTTCGCGAACACTAGGGTTTGGGTCTTCGAGAAGACCTATTCTGGCTTTT | Forward for lexA-4 gRNA |
| lexA-4R | AGGAAAAGCCAGAATAGGTCTTCTCGAAGACCCAAACCCTAGTGTTCGCGAAATTGG | Reverse for lexA-4 gRNA |
| ligA-1F | AAACCACAACTTGGACAATGGGTGTTTGGGTCTTCGAGAAGACCTATTCGTCTGGAC | Forward for ligA-1 gRNA |
| ligA-1R | AGGGTCCAGACGAATAGGTCTTCTCGAAGACCCAAACACCCATTGTCCAAGTTGTGG | Reverse for ligA-1 gRNA |
| ligA-2F | AAAACACGTGGTGATGGAACAACGTTTGGGTCTTCGAGAAGACCTATTCTGTTGATG | Forward for ligA-2 gRNA |
| ligA-2R | AGGCATCAACAGAATAGGTCTTCTCGAAGACCCAAACGTTGTTCCATCACCACGTGT | Reverse for ligA-2 gRNA |
| ligA-3F | AAATATTCCAGAACGTAGACCTGGTTTGGGTCTTCGAGAAGACCTATTCCGGAGAAC | Forward for ligA-3 gRNA |
| ligA-3R | AGGGTTCTCCGGAATAGGTCTTCTCGAAGACCCAAACCAGGTCTACGTTCTGGAATA | Reverse for ligA-3 gRNA |
| ligA-4F | AAAATTACTAACGGTAACTGAAGGTTTGGGTCTTCGAGAAGACCTATTCGCAACCGA | Forward for ligA-4 gRNA |

| | | |
|---|---|---|
| ligA-4R | AGGTCGGTTGCGAATAGGTCTTCTCGAAGAC CCAAACCTTCAGTTACCGTTAGTAAT | Reverse for ligA-4 gRNA |
| MT-1F | AAATTTCTACAGACAATGCAAGTGTTTGGGT CTTCGAGAAGACCTATTCTTGGGGTA | Forward for MT-1 gRNA |
| MT-1R | AGGTACCCCAAGAATAGGTCTTCTCGAAGA CCCAAACACTTGCATTGTCTGTAGAAA | Reverse for MT-1 gRNA |
| MT-2F | AAATTCTACAGACAATGCAAGTTGTTTGGGT CTTCGAGAAGACCTATTCTCACCAGC | Forward for MT-2 gRNA |
| MT-2R | AGGGCTGGTGAGAATAGGTCTTCTCGAAGA CCCAAACAACTTGCATTGTCTGTAGAA | Reverse for MT-2 gRNA |
| nadE-1F | AAAAGCGTATCTTGGTGCGCCAAGTTTGGGT CTTCGAGAAGACCTATTCGCTGTTTC | Forward for nadE-1 gRNA |
| nadE-1R | AGGGAAACAGCGAATAGGTCTTCTCGAAGA CCCAAACTTGGCGCACCAAGATACGCT | Reverse for nadE-1 gRNA |
| nadE-2F | AAAGTTAGGTATTTCAGGAGGACGTTTGGGT CTTCGAGAAGACCTATTCAACAGACG | Forward for nadE-2 gRNA |
| nadE-2R | AGGCGTCTGTTGAATAGGTCTTCTCGAAGAC CCAAACGTCCTCCTGAAATACCTAAC | Reverse for nadE-2 gRNA |
| nadE-3F | AAATTTGGTTTGAATAAACGACAGTTTGGGT CTTCGAGAAGACCTATTCGTTAAAGC | Forward for nadE-3 gRNA |
| nadE-3R | AGGGCTTTAACGAATAGGTCTTCTCGAAGAC CCAAACTGTCGTTTATTCAAACCAAA | Reverse for nadE-3 gRNA |
| nadE-4F | AAATTGCATATACAAGATACACGGTTTGGGT CTTCGAGAAGACCTATTCCCAGTTCT | Forward for nadE-4 gRNA |
| nadE-4R | AGGAGAACTGGGAATAGGTCTTCTCGAAGA CCCAAACCGTGTATCTTGTATATGCAA | Reverse for nadE-4 gRNA |
| NC1-1F | AAAGTTGGAACGATAGTTGGTGAGTTTGGGT CTTCGAGAAGACCTATTCCGCAAGGA | Forward for NC1-1 gRNA |
| NC1-1R | AGGTCCTTGCGGAATAGGTCTTCTCGAAGAC CCAAACTCACCAACTATCGTTCCAAC | Reverse for NC1-1 gRNA |
| NC2-2F | AAAAACGGACAAATAGTAGTCATGTTTGGG TCTTCGAGAAGACCTATTCAAGATTCG | Forward for NC2-2 gRNA |

| | | |
|---|---|---|
| NC2-2R | AGGCGAATCTTGAATAGGTCTTCTCGAAGAC<br>CCAAACATGACTACTATTTGTCCGTT | Reverse for NC2-2 gRNA |
| NT-2F | AAATGCTTGCCATAACATATCGAGTTTGGGT<br>CTTCGAGAAGACCTATTCGCGCGTAA | Forward for NT-2 gRNA |
| NT-2R | AGGTTACGCGCGAATAGGTCTTCTCGAAGAC<br>CCAAACTCGATATGTTATGGCAAGCA | Reverse for NT-2 gRNA |
| nusA-1F | AAAGAAGCTGTTGTTGAAGAGCTGTTTGGGT<br>CTTCGAGAAGACCTATTCACGTGCCA | Forward for nusA-1 gRNA |
| nusA-1R | AGGTGGCACGTGAATAGGTCTTCTCGAAGA<br>CCCAAACAGCTCTTCAACAACAGCTTC | Reverse for nusA-1 gRNA |
| nusA-2F | AAAGCTGTTGTTGAAGAGCTAGGGTTTGGGT<br>CTTCGAGAAGACCTATTCGACCTTAT | Forward for nusA-2 gRNA |
| nusA-2R | AGGATAAGGTCGAATAGGTCTTCTCGAAGA<br>CCCAAACCCTAGCTCTTCAACAACAGC | Reverse for nusA-2 gRNA |
| nusA-3F | AAAGATGCTGTTGGTGCATGTGTGTTTGGGT<br>CTTCGAGAAGACCTATTCTTCTGGGA | Forward for nusA-3 gRNA |
| nusA-3R | AGGTCCCAGAAGAATAGGTCTTCTCGAAGA<br>CCCAAACACACATGCACCAACAGCATC | Reverse for nusA-3 gRNA |
| nusA-4F | AAAATACATAACGATGGTCAACAGTTTGGG<br>TCTTCGAGAAGACCTATTCGAGAATAG | Forward for nusA-4 gRNA |
| nusA-4R | AGGCTATTCTCGAATAGGTCTTCTCGAAGAC<br>CCAAACTGTTGACCATCGTTATGTAT | Reverse for nusA-4 gRNA |
| recA-1F | AAACGAACGAATGGGTCAAGGTAGTTTGGG<br>TCTTCGAGAAGACCTATTCAGCACAGC | Forward for recA-1 gRNA |
| recA-1R | AGGGCTGTGCTGAATAGGTCTTCTCGAAGAC<br>CCAAACTACCTTGACCCATTCGTTCG | Reverse for recA-1 gRNA |
| recA-2F | AAAAGTACAAAGTAATGGCGGGGGTTTGGG<br>TCTTCGAGAAGACCTATTCTTTCCCAC | Forward for recA-2 gRNA |
| recA-2R | AGGGTGGGAAAGAATAGGTCTTCTCGAAGA<br>CCCAAACCCCCGCCATTACTTTGTACT | Reverse for recA-2 gRNA |
| recA-4F | AAAGAAATGGGAGACACTCACGTGTTTGGG<br>TCTTCGAGAAGACCTATTCGACTAGCT | Forward for recA-4 gRNA |

| | | |
|---|---|---|
| recA-4R | AGGAGCTAGTCGAATAGGTCTTCTCGAAGA CCCAAACACGTGAGTGTCTCCCATTTC | Reverse for recA-4 gRNA |
| recF-1F | AAACCTGACGTGAATATCCTCATGTTTGGGT CTTCGAGAAGACCTATTCCAGGAACC | Forward for recF-1 gRNA |
| recF-1R | AGGGGTTCCTGGAATAGGTCTTCTCGAAGAC CCAAACATGAGGATATTCACGTCAGG | Reverse for recF-1 gRNA |
| recF-2F | AAACCAATGAGGATATTCACGTCGTTTGGGT CTTCGAGAAGACCTATTCCCAGAGAA | Forward for recF-2 gRNA |
| recF-2R | AGGTTCTCTGGGAATAGGTCTTCTCGAAGAC CCAAACGACGTGAATATCCTCATTGG | Reverse for recF-2 gRNA |
| recF-3F | AAAGCGCAAATAGAACCACATTGGTTTGGG TCTTCGAGAAGACCTATTCCTCGCTTG | Forward for recF-3 gRNA |
| recF-3R | AGGCAAGCGAGGAATAGGTCTTCTCGAAGA CCCAAACCAATGTGGTTCTATTTGCGC | Reverse for recF-3 gRNA |
| recF-4F | AAAGGTGAGCTTAGTTATAGACAGTTTGGGT CTTCGAGAAGACCTATTCCAGAATGA | Forward for recF-4 gRNA |
| recF-4R | AGGTCATTCTGGAATAGGTCTTCTCGAAGAC CCAAACTGTCTATAACTAAGCTCACC | Reverse for recF-4 gRNA |
| rpmD-1F | AAAAAGATAACCCTGCTATTCGTGTTTGGGT CTTCGAGAAGACCTATTCCTGACAGC | Forward for rpmD-1 gRNA |
| rpmD-1R | AGGGCTGTCAGGAATAGGTCTTCTCGAAGA CCCAAACACGAATAGCAGGGTTATCTT | Reverse for rpmD-1 gRNA |
| rpmD-2F | AAACGTAAAACTGTTGAAGCTTTGTTTGGGT CTTCGAGAAGACCTATTCTGCTTTGT | Forward for rpmD-2 gRNA |
| rpmD-2R | AGGACAAAGCAGAATAGGTCTTCTCGAAGA CCCAAACAAAGCTTCAACAGTTTTACG | Reverse for rpmD-2 gRNA |
| rpmD-3F | AAAGACCAATAACACTACGAGTGGTTTGGG TCTTCGAGAAGACCTATTCAAAGGCCG | Forward for rpmD-3 gRNA |
| rpmD-3R | AGGCGGCCTTTGAATAGGTCTTCTCGAAGAC CCAAACCACTCGTAGTGTTATTGGTC | Reverse for rpmD-3 gRNA |
| rpmD-4F | AAATGATTTGCCCACGAATAGCAGTTTGGGT CTTCGAGAAGACCTATTCGGCAATAG | Forward for rpmD-4 gRNA |

| | | |
|---|---|---|
| rpmD-4R | AGGCTATTGCCGAATAGGTCTTCTCGAAGAC CCAAACTGCTATTCGTGGGCAAATCA | Reverse for rpmD-4 gRNA |
| rpoB-1F | AAATTGGGACGGTTACAACTATGGTTTGGGT CTTCGAGAAGACCTATTCTTTGGAAG | Forward for rpoB-1 gRNA |
| rpoB-1R | AGGCTTCCAAAGAATAGGTCTTCTCGAAGAC CCAAACCATAGTTGTAACCGTCCCAA | Reverse for rpoB-1 gRNA |
| rpoB-2F | AAATTTTGGTGAGATGGAGGTATGTTTGGGT CTTCGAGAAGACCTATTCTCAGCAAT | Forward for rpoB-2 gRNA |
| rpoB-2R | AGGATTGCTGAGAATAGGTCTTCTCGAAGAC CCAAACATACCTCCATCTCACCAAAA | Reverse for rpoB-2 gRNA |
| rpoB-3F | AAAAGGACGTGTGAAAACATACGGTTTGGG TCTTCGAGAAGACCTATTCTACAGCTC | Forward for rpoB-3 gRNA |
| rpoB-3R | AGGGAGCTGTAGAATAGGTCTTCTCGAAGA CCCAAACCGTATGTTTTCACACGTCCT | Reverse for rpoB-3 gRNA |
| rpoB-4F | AAAGCTATTACAGCTAAGCACAGGTTTGGGT CTTCGAGAAGACCTATTCACTTCTGT | Forward for rpoB-4 gRNA |
| rpoB-4R | AGGACAGAAGTGAATAGGTCTTCTCGAAGA CCCAAACCTGTGCTTAGCTGTAATAGC | Reverse for rpoB-4 gRNA |
| rpsL-1F | AAAGTAAGTTATGTCCGATACCAGTTTGGGT CTTCGAGAAGACCTATTCCAAATGGA | Forward for rpsL-1 gRNA |
| rpsL-1R | AGGTCCATTTGGAATAGGTCTTCTCGAAGAC CCAAACTGGTATCGGACATAACTTAC | Reverse for rpsL-1 gRNA |
| rpsL-2F | AAAAGTGTTGTACTTGTACGTGGGTTTGGGT CTTCGAGAAGACCTATTCTGGAAATC | Forward for rpsL-2 gRNA |
| rpsL-2R | AGGGATTTCCAGAATAGGTCTTCTCGAAGAC CCAAACCCACGTACAAGTACAACACT | Reverse for rpsL-2 gRNA |
| rpsL-3F | AAACACAGTGTTGTACTTGTACGGTTTGGGT CTTCGAGAAGACCTATTCTAGCTGGA | Forward for rpsL-3 gRNA |
| rpsL-3R | AGGTCCAGCTAGAATAGGTCTTCTCGAAGAC CCAAACCGTACAAGTACAACACTGTG | Reverse for rpsL-3 gRNA |
| rpsL-4F | AAATTAAACTCACCACAAAAACGGTTTGGG TCTTCGAGAAGACCTATTCTGGCGAGT | Forward for rpsL-4 gRNA |

| | | |
|---|---|---|
| rpsL-4R | AGGACTCGCCAGAATAGGTCTTCTCGAAGA CCCAAACCGTTTTTGTGGTGAGTTTAA | Reverse for rpsL-4 gRNA |
| secY-1F | AAATTGGACAAACTGCGTTCGTTGTTTGGGT CTTCGAGAAGACCTATTCTCTCAGCG | Forward for secY-1 gRNA |
| secY-1R | AGGCGCTGAGAGAATAGGTCTTCTCGAAGA CCCAAACAACGAACGCAGTTTGTCCAA | Reverse for secY-1 gRNA |
| secY-2F | AAATGGGCAAAACAAGGTGAAGTGTTTGGG TCTTCGAGAAGACCTATTCTTAAACGC | Forward for secY-2 gRNA |
| secY-2R | AGGGCGTTTAAGAATAGGTCTTCTCGAAGAC CCAAACACTTCACCTTGTTTTGCCCA | Reverse for secY-2 gRNA |
| secY-3F | AAAAAAAGTATTTAATAACTCAGGTTTGGGT CTTCGAGAAGACCTATTCCGAAACCC | Forward for secY-3 gRNA |
| secY-3R | AGGGGGTTTCGGAATAGGTCTTCTCGAAGAC CCAAACCTGAGTTATTAAATACTTTT | Reverse for secY-3 gRNA |
| secY-4F | AAAAAAGATTACAGGAATAACACGTTTGGG TCTTCGAGAAGACCTATTCGGGTGCTT | Forward for secY-4 gRNA |
| secY-4R | AGGAAGCACCCGAATAGGTCTTCTCGAAGA CCCAAACGTGTTATTCCTGTAATCTTT | Reverse for secY-4 gRNA |
| sigB-1F | AAAGTTGGTATGGTTGGTTTAATGTTTGGGT CTTCGAGAAGACCTATTCCACAAACA | Forward for sigB-1 gRNA |
| sigB-1R | AGGTGTTTGTGGAATAGGTCTTCTCGAAGAC CCAAACATTAAACCAACCATACCAAC | Reverse for sigB-1 gRNA |
| sigB-2F | AAAAAAGAGACAGGTGAGCGTATGTTTGGG TCTTCGAGAAGACCTATTCTCAGAACG | Forward for sigB-2 gRNA |
| sigB-2R | AGGCGTTCTGAGAATAGGTCTTCTCGAAGAC CCAAACATACGCTCACCTGTCTCTTT | Reverse for sigB-2 gRNA |
| sigB-3F | AAAAGAAGAAGTGTTAGAAGCAAGTTTGGG TCTTCGAGAAGACCTATTCACGTATTG | Forward for sigB-3 gRNA |
| sigB-3R | AGGCAATACGTGAATAGGTCTTCTCGAAGA CCCAAACTTGCTTCTAACACTTCTTCT | Reverse for sigB-3 gRNA |
| sigB-4F | AAAAGGTGAACGCTCTAATTCAGGTTTGGGT CTTCGAGAAGACCTATTCCCTAATAC | Forward for sigB-4 gRNA |

| | | |
|---|---|---|
| sigB-4R | AGGGTATTAGGGAATAGGTCTTCTCGAAGA CCCAAACCTGAATTAGAGCGTTCACCT | Reverse for sigB-4 gRNA |
| sosA-1F | AAAACAATAGCGAACAAAGAGATGTTTGGG TCTTCGAGAAGACCTATTCTGGCGCTT | Forward for sosA-1 gRNA |
| sosA-1R | AGGAAGCGCCAGAATAGGTCTTCTCGAAGA CCCAAACATCTCTTTGTTCGCTATTGT | Reverse for sosA-1 gRNA |
| sosA-2F | AAATTTGTTCGCTATTGTTTGTAGTTTGGGTC TTCGAGAAGACCTATTCCTTGATCC | Forward for sosA-2 gRNA |
| sosA-2R | AGGGGATCAAGGAATAGGTCTTCTCGAAGA CCCAAACTACAAACAATAGCGAACAAA | Reverse for sosA-2 gRNA |
| sosA-3F | AAATGCGAACATTAGTGCTCACTGTTTGGGT CTTCGAGAAGACCTATTCTAACTTGG | Forward for sosA-3 gRNA |
| sosA-3R | AGGCCAAGTTAGAATAGGTCTTCTCGAAGA CCCAAACAGTGAGCACTAATGTTCGCA | Reverse for sosA-3 gRNA |
| sosA-4F | AAAACTTAACAAGAATATTGCAAGTTTGGGT CTTCGAGAAGACCTATTCACTACCCT | Forward for sosA-4 gRNA |
| sosA-4R | AGGAGGGTAGTGAATAGGTCTTCTCGAAGA CCCAAACTTGCAATATTCTTGTTAAGT | Reverse for sosA-4 gRNA |
| trmD-1F | AAAATACCTTGTCCGCCACCATAGTTTGGGT CTTCGAGAAGACCTATTCACGGGTTA | Forward for trmD-1 gRNA |
| trmD-1R | AGGTAACCCGTGAATAGGTCTTCTCGAAGAC CCAAACTATGGTGGCGGACAAGGTAT | Reverse for trmD-1 gRNA |
| trmD-2F | AAAGTATGGTGGCGGACAAGGTAGTTTGGG TCTTCGAGAAGACCTATTCGAGATAGA | Forward for trmD-2 gRNA |
| trmD-2R | AGGTCTATCTCGAATAGGTCTTCTCGAAGAC CCAAACTACCTTGTCCGCCACCATAC | Reverse for trmD-2 gRNA |
| trmD-3F | AAAGTTAGACCCTTAAATTCACGGTTTGGGT CTTCGAGAAGACCTATTCTGTTCATG | Forward for trmD-3 gRNA |
| trmD-3R | AGGCATGAACAGAATAGGTCTTCTCGAAGA CCCAAACCGTGAATTTAAGGGTCTAAC | Reverse for trmD-3 gRNA |
| trmD-4F | AAAGTCCGCATATGAAAACGATGGTTTGGG TCTTCGAGAAGACCTATTCTCACTGAT | Forward for trmD-4 gRNA |

| | | |
|---|---|---|
| trmD-4R | AGGATCAGTGAGAATAGGTCTTCTCGAAGA CCCAAACCATCGTTTTCATATGCGGAC | Reverse for trmD-4 gRNA |
| umuC-1F | AAATTTGGCTATAGTGATGAAGGGTTTGGGT CTTCGAGAAGACCTATTCGTGATAAC | Forward for umuC-1 gRNA |
| umuC-1R | AGGGTTATCACGAATAGGTCTTCTCGAAGAC CCAAACCCTTCATCACTATAGCCAAA | Reverse for umuC-1 gRNA |
| umuC-2F | AAAGTGTTTCTTGTATTGAAAAGGTTTGGGT CTTCGAGAAGACCTATTCGCTAAGGC | Forward for umuC-2 gRNA |
| umuC-2R | AGGGCCTTAGCGAATAGGTCTTCTCGAAGAC CCAAACCTTTTCAATACAAGAAACAC | Reverse for umuC-2 gRNA |
| umuC-3F | AAAGCGTCAGGGTTCTGTAATATGTTTGGGT CTTCGAGAAGACCTATTCCTCTAACT | Forward for umuC-3 gRNA |
| umuC-3R | AGGAGTTAGAGGAATAGGTCTTCTCGAAGA CCCAAACATATTACAGAACCCTGACGC | Reverse for umuC-3 gRNA |
| umuC-4F | AAAGCCTTCATCACTATAGCCAAGTTTGGGT CTTCGAGAAGACCTATTCATACGGGG | Forward for umuC-4 gRNA |
| umuC-4R | AGGCCCCGTATGAATAGGTCTTCTCGAAGAC CCAAACTTGGCTATAGTGATGAAGGC | Reverse for umuC-4 gRNA |
